# HIV–1 infection reduces NAD capping of host cell snRNA and snoRNA

**DOI:** 10.1101/2022.11.10.515957

**Authors:** Barbora Benoni, Jiří František Potužník, Anton Škríba, Roberto Benoni, Jana Trylcova, Matouš Tulpa, Kristína Spustová, Katarzyna Grab, Maria-Bianca Mititelu, Jan Pačes, Jan Weber, David Stanek, Joanna Kowalska, Lucie Bednarova, Zuzana Keckesova, Pavel Vopalensky, Lenka Gahurova, Hana Cahova

## Abstract

Nicotinamide adenine dinucleotide (NAD) is a critical component of the cellular metabolism and also serves as an alternative 5′ cap on various RNAs. However, the function of the NAD RNA cap is still under investigation. We studied NAD capping of RNAs in HIV–1–infected cells because HIV–1 is responsible for the depletion of the NAD/NADH cellular pool and causing intracellular pellagra. By applying the NAD captureSeq protocol to HIV–1–infected and uninfected cells, we revealed that four snRNAs (e.g. U1) and four snoRNAs lost their NAD cap when infected with HIV–1. Here, we provide evidence that the presence of the NAD cap decreases the stability of the U1/HIV–1 pre–mRNA duplex. Additionally, we demonstrate that reducing the quantity of NAD–capped RNA by overexpressing the NAD RNA decapping enzyme DXO results in an increase in HIV–1 infectivity. This suggests that NAD capping is unfavorable for HIV–1 and plays a role in its infectivity.

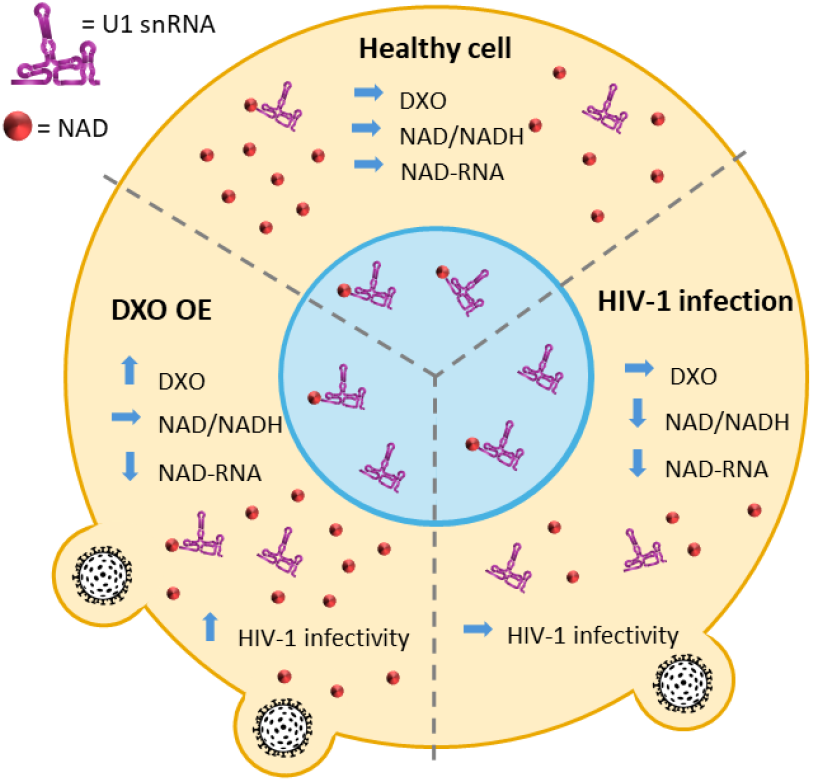

---

To date, more than 170 RNA modifications have been discovered^*1*^. Among the least explored RNA modifications are 5′ non–canonical RNA caps, including Nicotinamide adenine dinucleotide (NAD)^*2*^, Flavin adenine dinucleotide^*3*^ and dinucleoside polyphosphates^*4, 5*^. Because they are typically detected through LC–MS analysis of digested RNA, there is no information on which RNA sequences bear these caps. Several methods have been developed for the sequencing of NAD–RNA. The original NAD captureSeq protocol ^*6, 7*^ was later modified, resulting in various protocols such as SPAAC–NAD–Seq^*8*^, or NADcapPro, circNC ^*9*^ and ONE–seq ^*10*^. These methods allow for the identification of NAD–RNA sequences in various organisms (e.g. *Staphylococcus aureus* ^*11*^, *Saccharomyces cerevisiae*^*12*^, human cells^*13*^). NAD has been detected as a 5′ RNA cap attached to small regulatory RNAs in bacteria^*6*^ and various mRNAs in higher organisms^*14*^. Recently, NAD–RNA was also reported to function as a substrate for the RNAylation of proteins upon phage infection in bacteria^*15*^. However, the role of this cap in higher organisms remains unclear.

Because HIV–1 infection of human cells depletes the cellular pool of free NAD^*16*^, we envisaged it as a physiologically interesting model system in which to study the role of the NAD RNA cap. There are two different mechanisms responsible for NAD depletion: First, there is an increased activity of CD38 in HIV– 1–infected cells, which reduces the NAD pool^*17*^. The second mechanism includes the activation of poly (ADP–ribose) polymerases (PARPs), induced by oxidative stress during HIV–1 infection,^*18, 19*^ which consume NAD and thus trigger *de novo* niacin synthesis (precursor of NAD). Moreover, it has been reported that nicotinamide acts as an inhibitor of HIV–1 infection^*20*^. For this reason, niacin has been suggested as a potential AIDS preventive factor^*21*^. In addition, the genetic variation in the locus of the NAD decapping enzyme DXO is associated with differences in the response to HIV–1 infection^*22*^.

Here, we report that subsets of cellular snRNAs and snoRNAs lose the NAD cap after HIV–1 infection. Among the identified RNAs that lose their NAD cap upon HIV–1 infection, we detected U1 snRNA, which is essential for viral replication^*23, 24*^. We found that in comparison with 7,7,2–trimethylguanosine (TMG)– capped U1 snRNA^*25*^, the NAD cap has a destabilizing effect on the binding of U1 snRNA to viral pre–mRNA. This suggests that the NAD RNA cap reduces the binding of U1 snRNA with the viral RNA target and might thus contribute to the protection against HIV–1 infection. To test this hypothesis, we overexpressed and knocked down the NAD decapping enzyme DXO and monitored HIV–1 production. The overexpression of DXO, which decreases cellular levels of NAD–capped RNAs, results in higher virus infectivity. These findings provide the first link between cellular NAD/NADH levels, NAD capping, and HIV–1 infectivity.

It has been reported that HIV–1 infection causes intracellular pellagra, meaning the depletion of the NAD/NADH pool in the cell^*16*^; therefore, we measured the total concentration of NAD in control (uninfected) MT4 cells and MT4 cells infected with HIV–1 (Figure 1A). Indeed, we observed a nearly four times greater intracellular concentration of NAD in control cells compared to infected cells. Because NAD is incorporated into RNA co–transcriptionally by RNA polymerases as a non–canonical initiating nucleotide^*26*^, we hypothesized that a change in the intracellular level of NAD may also influence the NAD capping of RNA.

**Figure 1.**
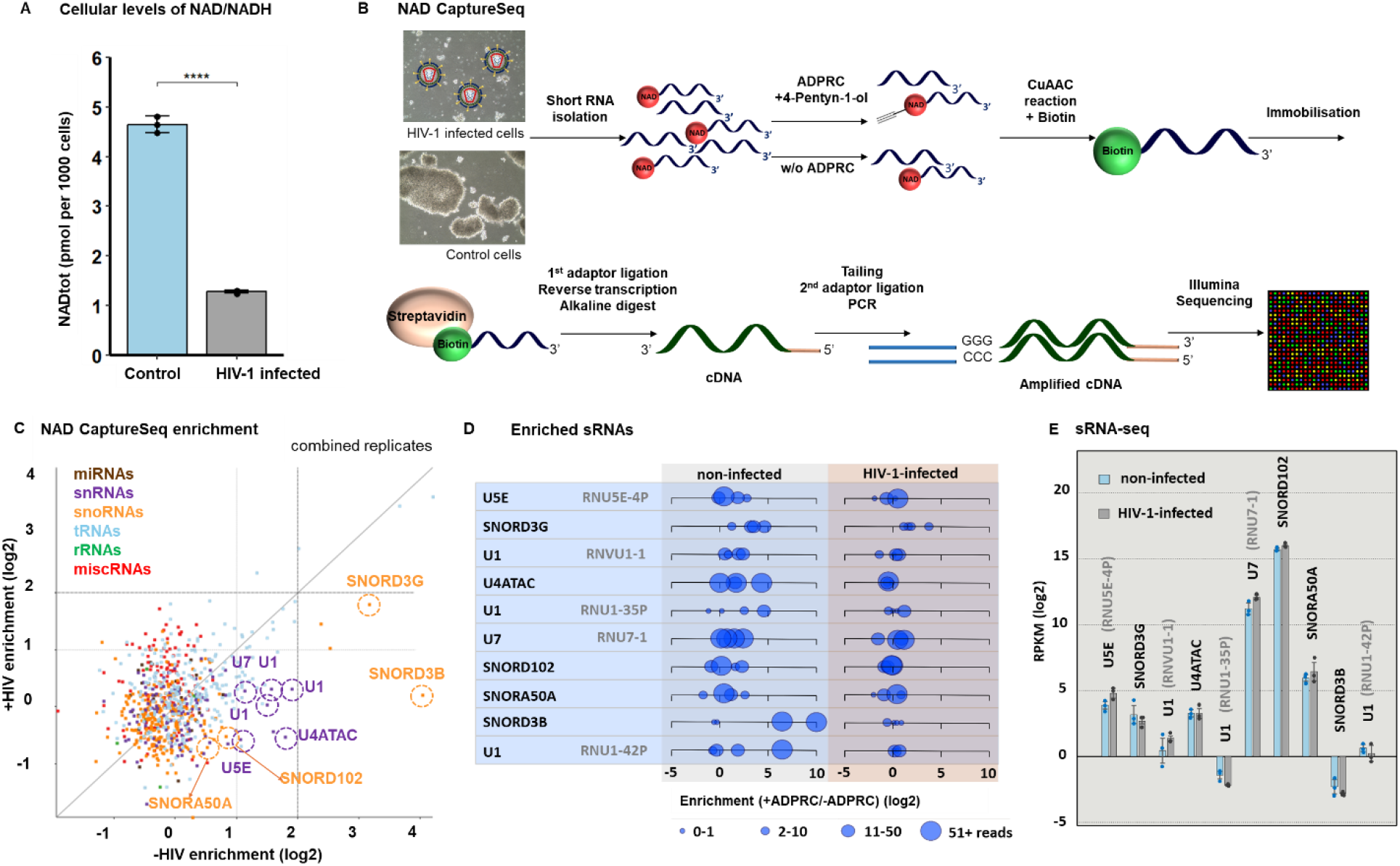
Changes in NAD RNA capping upon HIV–1 infection. (A) Levels of free NAD/NADH in control and HIV–1–infected cells (measured in biological triplicates and technical duplicates, t–test). (B) Scheme of the NAD captureSeq protocol applied to control and HIV–1–infected cells. (C) Scatter plot showing the NAD captureSeq enrichment (+ADPRC vs −ADPRC) in control (X axis) and HIV–1–infected samples (Y axis), showing average values across replicates. Highlighted sRNAs passed the criteria listed in FigS1B, i.e. change in NAD cap abundance upon HIV–1 infection, having specific enrichment in +ADPRC samples, sufficient read count and consistency across replicates; these sRNAs are visualized in more detail for individual replicates in Fig1D. (D) Enrichment of RNAs in NAD captureSeq analysis enriched in control cells versus HIV–1 infected cells, matching highlighted sRNAs in Fig1C. Blue circles show individual replicates, size of the circle corresponds to the number of mapped reads. (E) sRNA–seq analysis of identified candidate sRNAs in control and HIV–1–infected cells (prepared in biological triplicates).

As studies of the NAD cap in mRNA are yet to arrive at a clear–cut explanation of its role, we focused on the potential effect of the NAD RNA cap on short non–coding regulatory RNAs (sRNA). The fraction of sRNA usually contains various regulatory RNAs which bind to their targets and where RNA caps could affect the recognition and binding and thus it might be easier to study the functional role of the cap itself. We therefore investigated how HIV–1 infection affects the extent of NAD capping of sRNA. To identify RNAs with altered NAD capping, we prepared four NAD captureSeq libraries from the sRNA fraction of control and HIV–1–infected cells. The NAD captureSeq protocol^6, 7^ relies on a selective reaction of ADP– ribosyl cyclase (ADPRC) with 4–pentyn–1–ol and with NAD–RNA (Figure 1B, Table S1), which allows for the enrichment of previously NAD–capped RNA and subsequent RNAseq analysis. To sort out non– specifically interacting RNA, ADPRC was omitted in negative control samples (–ADPRC). Even though the m^7^G cap has been reported to partially react with ADPRC^*9*^, the sRNA fraction we used contained approximately 5 times less m^7^Gp_3_Am (cap1) per molecule of RNA than the long RNA (lRNA) fraction (Figure S1A). We therefore did not deplete m^7^G–capped RNA from our sRNA.

The sequenced NAD captureSeq libraries were mapped to both the human genome and that of HIV–1 (Figure S1B). We detected the NAD cap in 94 cellular sRNAs (2 miRNAs, 6 miscellaneous RNAs, abbreviated to miscRNA, 62 tRNAs, 12 snoRNAs, 12 snRNAs). In line with previous observations^*13*^, we also detected the NAD cap on the 5′–end fragments of some mRNAs but not on the 5′ end of viral mRNA fragments (Figure S1C), which was previously shown to contain the TMG or m^7^G cap^*27*^. Overall, we observed similar quantities of enriched and depleted NAD–capped sRNAs in both HIV–1–infected and control cells (Figure 1C). However, with 42 NAD–capped sRNAs, these quantities varied substantially between HIV–1– infected and control cells. An increase in NAD capping upon HIV–1 infection was detected on sRNAs known in literature to be packed in HIV–1 virions (e.g. various tRNAs) (Figure S2) ^*28, 29*^ however we did not pursue this observation further. Particularly prominent were hits with decreased NAD capping upon HIV–1 infection from two classes of sRNAs: snRNAs and snoRNAs. To filter out sRNAs that react non–specifically in the +ADPRC reaction (measured by low enrichment in +ADPRC vs −ADPRC samples), we applied a set of criteria regarding the number of mapped reads and consistency across replicates (detailed parameters are listed in Figure S1B, sRNAs that passed each filtering step are in Table S2 Supplemental Excel Table). After this filtration, we identified four snoRNAs (SNORD3G, SNORD102, SNORA50A and SNORD3B) and four snRNAs (U1, U4ATAC, U5E and U7) with decreased NAD capping upon HIV–1 infection (Figure 1C, D).

Because the amount of total RNA in control and HIV–1–infected cells was similar (Figure S3A), we further investigated whether the decreased amount of the NAD–cap on particular sRNAs after HIV–1 infection is caused by altered sRNA expression. We prepared sRNA–seq libraries from control and infected cells and performed DESeq2 analysis. Out of the 2014 detected sRNAs (miRNAs, miscRNAs, tRNAs, snRNAs, snoRNAs and rRNAs), 89 were identified as significantly differentially expressed: 52 sRNAs were upregulated and 37 downregulated after infection (Table S3, Figure S3B, C). Neither of these 89 sRNAs was enriched in the NAD captureSeq protocol. Moreover, using these data, we confirmed that the expression levels of identified snRNA and snoRNA do not change after HIV–1 infection (Figure 1E). The sequencing results were independently confirmed by RT–qPCR of selected sRNAs (Figure S4). These experiments exclude the possibility that changes in NAD–RNA enrichment in infected cells are caused by the differential expression of particular cellular RNAs upon HIV–1 infection.

Next, we also wanted to confirm the presence of the NAD cap in RNA from MT4 cells by means of LC–MS analysis. To avoid the detection of NAD non–covalently bound to RNA, we included a urea wash step^*30*^ in our protocol. Isolated and washed RNA was treated with the NudC enzyme to cleave nicotinamide mononucleotide (NMN) from NAD–capped RNA. The NMN was then converted to nicotinamide riboside (NR) by employing shrimp alkaline phosphatase^*30*^. For the purposes of quantification, isotopically labelled (deuterated) D_3_–NR was spiked in each sample as an internal standard. In general, we did not observe any statistical difference between infected and control cells in either sRNA or lRNA (Figure S5 A–E). This observation might be explained by the fact that similar quantities of sRNAs are enriched or depleted in NAD captureSeq data in HIV–1–infected and control cells (Figure 1C). On the other hand, LC–MS analysis of U1 RNA pulled down from the sRNA fraction isolated from control and HIV–1–infected cells confirmed the significant depletion of the NAD cap on this particular RNA (Figure S6 A–C). This finding shows that HIV–1 infection causes a decrease in the NAD capping of U1.

The U1 snRNA is known to play an important role in the lifecycle of HIV–1^*24, 31*^. The 5′ region (8 nucleotides) of U1 binds complementarily to the 5′ splice site of unspliced HIV–1 mRNA, which leads to the Rev– regulated translocation of partially spliced or unspliced HIV–1 mRNA into the cytosol^*23*^. A single mutation in the 5′ splice site of HIV–1 mRNA causes a significant decrease in HIV–1 replication due to limited binding of U1. This effect could be suppressed by co–transfection with U1 that restores base pairing with the mutated 5′ splice site of HIV–1 pre–mRNA, which indicates the importance of U1/HIV–1 pre–mRNA base pairing^*23*^. We hypothesized that altered RNA capping of U1 (the 5’ end that recognizes HIV–1 pre–mRNA) might influence the binding to pre–mRNA and its subsequent interaction with the Rev protein. This complex is then translocated to the cytosol and leads to the translation of envelope–encoding HIV–1 mRNA. In this way, the alteration of U1 capping might influence the viral replication cycle. For these reasons, we investigated the role of the 5′ RNA cap of U1 in binding to HIV–1 pre–mRNA. Until now, it was assumed that U1 is naturally capped only with the TMG cap^*25*^. Here, we show that part of the cellular pool of U1 is also capped with NAD and that this cap gets depleted upon HIV–1 infection. We recently showed that some non–canonical RNA caps (e.g. Ap_3_G) bind to DNA templates via non–canonical base pairing during transcription^*32*^. Accordingly, we hypothesize that the NAD RNA cap may contribute to the binding of U1 to its target HIV–1 pre–mRNA and that it may negatively or positively influence the strength of the interaction. To test this hypothesis, we prepared 20–mer RNA mimicking the 5′ U1 RNA region with either the NAD or the TMG cap (Supplementary protocol) through *in vitro* transcription with T7 RNA polymerase (the original sequence AUA was changed to AGG due to the template requirements of T7 RNA polymerase). Pseudouridine triphosphate (ΨTP) was used instead of UTP to mimic the natural presence of Ψ in positions 5 and 6 in U1 (Figure 2A). Further, we prepared 20–mer RNA mimicking the D4 splice site of HIV–1 pre–mRNA complementary to U1. We measured the melting temperature (*T*_m_) values of corresponding duplexes using an intercalating fluorophore and a light cycler device. The *T*_m_ of RNA duplexes in this assay were 70.5 ± 0.6°C and 64.6 ± 0.8°C for TMG–U1 and NAD–U1, respectively (Figure 2B). As *in vitro* transcription with ΨTP did not lead to a sufficient amount of RNA for circular dichroism spectroscopy (CD) studies, which are usually used for RNA duplex characterization, we prepared a 20–mer mimicking the 5′ U1 RNA region with either the NAD or TMG cap without ΨTP (Figure S7A). The CD spectra of both duplexes with a positive maximum at 264 nm were typical for the right–handed A–type double helix of RNA (Figure S7B,C) and confirmed the observation from the melting curve analysis on the light cycler device (Figure S7D). The difference in the stability of duplexes with and without pseudouridine may be explained by a certain rigidity caused by the pseudouridines, which restricts the motion of neighbouring nucleotides including the 5′ cap^*33, 34*^. This experiment showed that the binding of TMG–U1 to HIV–1 pre–mRNA is much stronger than the NAD–U1 interaction with HIV–1 pre–mRNA. This indicates that the NAD cap reduces the binding of U1 to the target HIV–1 pre–mRNA sequence and thus may negatively influence HIV–1 replication.

**Figure 2.**
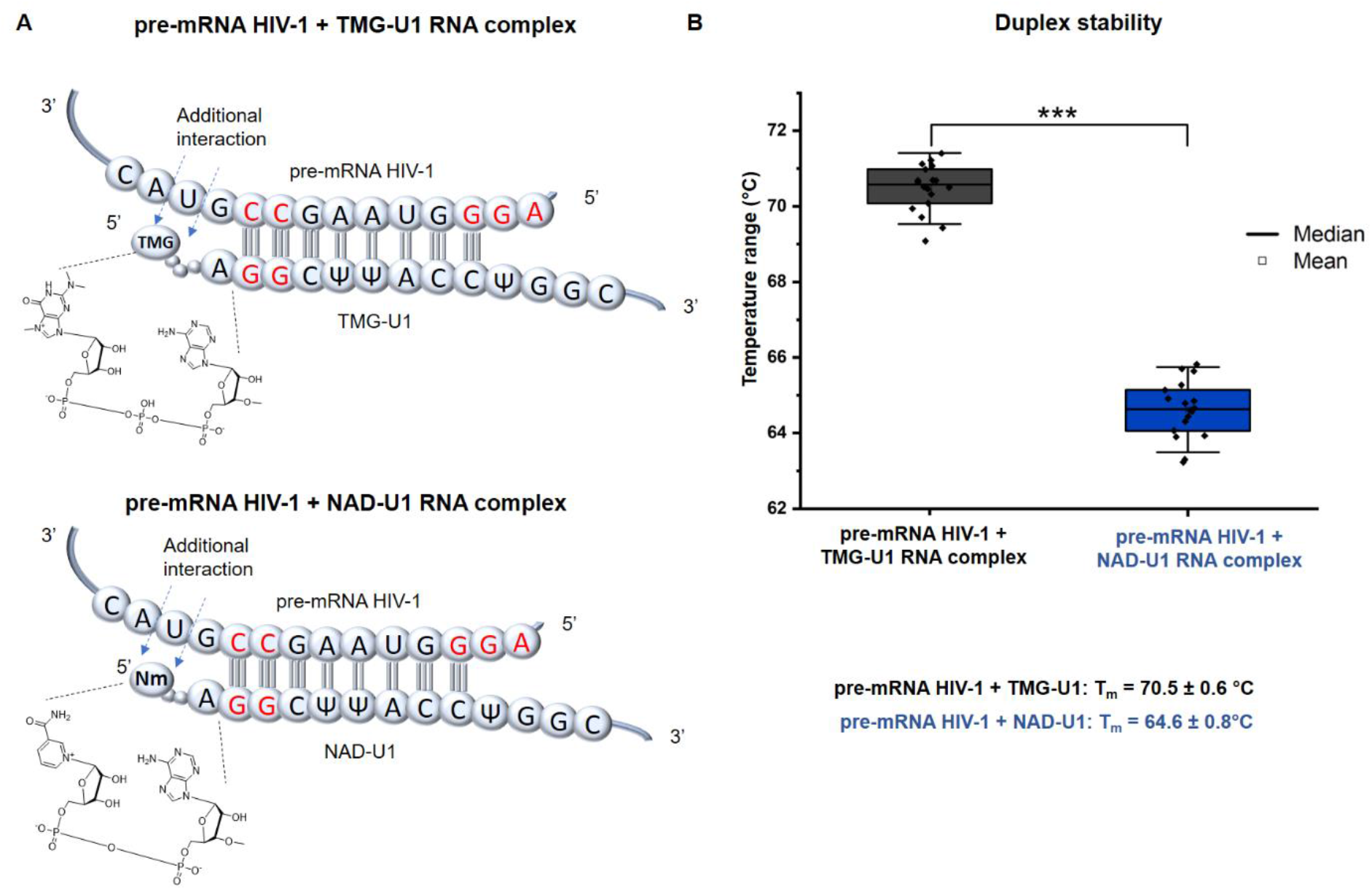
NAD cap of U1 snRNA destabilizes the complex with HIV–1 pre–mRNA. (A) Scheme of the complex formed by complemental regions of pre–mRNA HIV–1 with either TMG–U1 (left) or NAD–U1 (right). Nucleobase symbols in red mark positions mutated to meet the requirements of T7 RNA polymerase. (B) Duplex stability of pre–mRNA HIV–1 with TMG–U1 (dark grey) or NAD–U1 (blue) measured using a light cycler device (measured in triplicate in series of nine measurements of each, t–test).

The NAD cap is removed in human cells by these enzymes: Nudt12 ^*35*^, Nudt16^*36*^ and DXO^*13*^. Nudt12 and Nudt16 members of the NudiX family, cleave the pyrophosphate backbone of free NAD^*37*^ and NAD–RNA and thus reduce the intracellular concentrations of free NAD and NAD–RNA whereas DXO cleaves the entire moiety of NAD from RNA and does not affect the intracellular concentration of free NAD (Figure 3A)^*13*^. To investigate solely the role of NAD–RNA (not free NAD) in HIV–1 infection, we focused on the DXO decapping enzyme. In the following experiments, we wanted to explore the direct effect of changed NAD RNA capping on HIV–1 infection and to distinguish it from the effect of the changed total NAD pool. Although it has been reported that DXO may cleave the TMG cap^*38*^, we show that NAD–RNA is its preferred substrate (Figure 3B, C). Next, we checked whether the depletion of the NAD cap on U1 RNA in HIV–1–infected cells is not caused by the upregulation of DXO or Nudt12 upon infection; however, Western blot and RT–qPCR analysis showed that the levels of DXO are similar in control and HIV–1– infected cells (Figure S8A, B). The Nudt12 was not detectable in any of these samples by RT–qPCR (Figure S8B).

**Figure 3.**
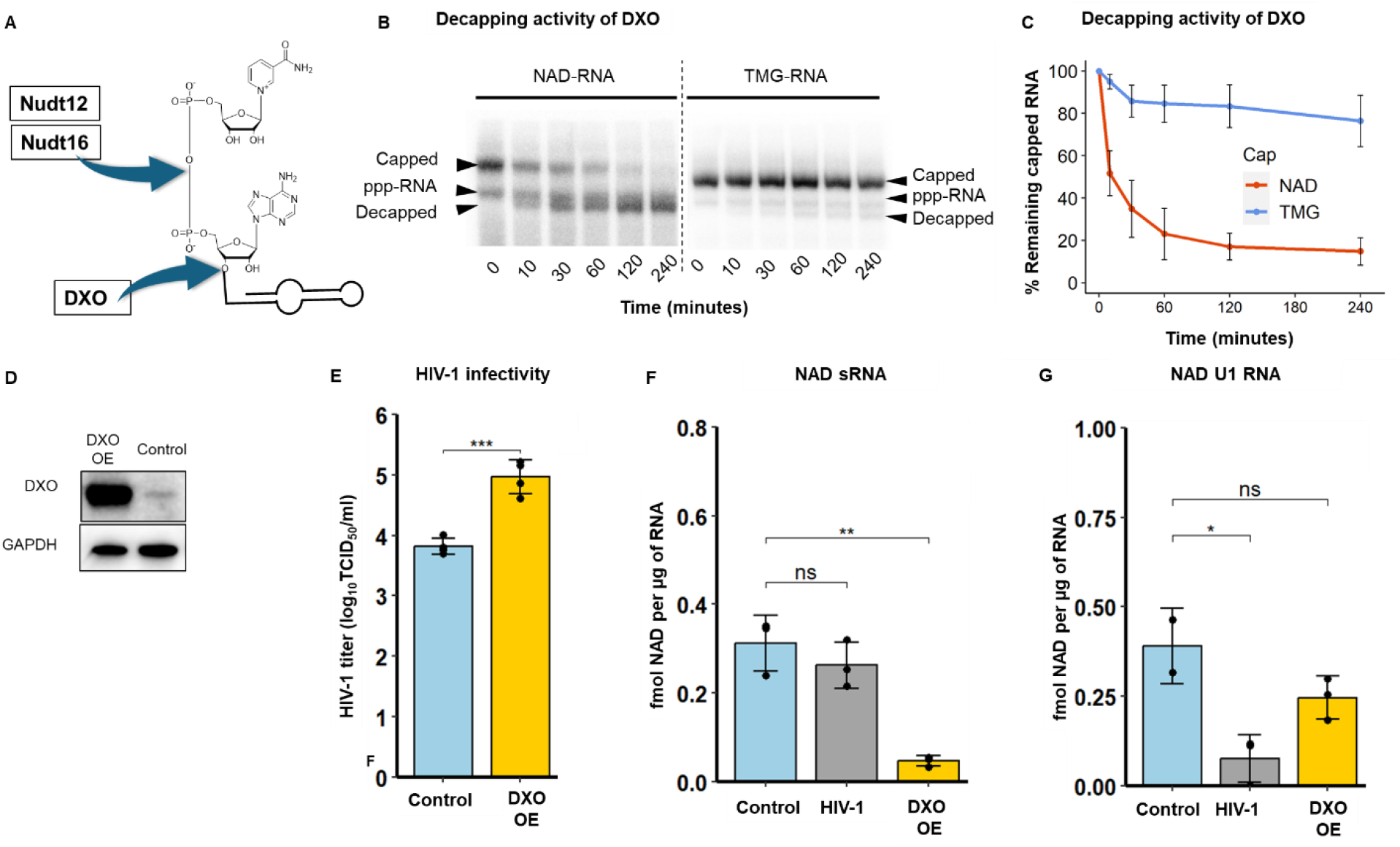
Decrease of NAD RNA capping leads to increased HIV–1 infectivity. (A) Cleavage positions of the NAD RNA cap by the two known decapping enzymes Nudt12 and DXO. (B) In vitro DXO cleavage of NAD–U1 and TMG–U1 at various concentrations of DXO (5 nM, 10 nM, 25 nM and 50 nM. (C) Quantification of DXO decapping activity based on PAGE gels (experiments performed in triplicate, error bar=standard deviation of the mean). (D) Western blot analysis of DXO and GAPDH (as a control) in control cells and cells with overexpressed (DXO OE). (E) HIV–1 infectivity determined in control cells and cells with overexpressed DXO (DXO OE) (ANOVA test). (F) LC–MS quantification of the NAD RNA cap level in sRNA isolated from control cells and cells with overexpressed DXO (DXO OE) (t–test). (G) LC–MS quantification of the NAD RNA cap level in pulled down U1 RNA (t–test). All the experiments were performed in biological triplicates.

Using lentiviral vectors, we prepared stable MT4 cell lines in which the DXO gene was either overexpressed (MT4–DXO OE) or downregulated (MT4–DXO KD). Indeed, overexpression of DXO led to a substantial increase in the levels of the enzymes in MT4–DXO OE cells (Figure 3D, Figure S9A) whereas DXO was almost completely diminished in MT4–DXO KD cells.

Further, we infected MT4–DXO OE and MT4–DXO KD cells with HIV–1, and after four days, we evaluated the effect of DXO expression on HIV–1 infectivity by determining the titer of HIV–1 (Figure 3E). The infectivity of HIV–1 in MT4–DXO KD cells was comparable to that in control MT4 cells (Figure S9B). By contrast, the overexpression of the NAD–RNA decapping enzyme DXO led to an increase in the infectivity of HIV–1 in comparison with control cells (Figure 3E).

We also followed the effect of DXO overexpression and knockdown on the level of NAD RNA capping in the sRNA fraction (Figure 3F and Figure S9C). We quantified the extent of NAD RNA capping in MT4–DXO OE and MT4–DXO KD cells by LC–MS (Figure S10A–C). Surprisingly, however, the extent of NAD capping of RNA isolated from MT4–DXO KD cells was lower than in control cells. One possible explanation might reside in the redundancy of decapping enzymes in general, which was previously described^*39, 40*^. To support this hypothesis, we performed RT–qPCR of NudiX enzymes known to cleave NAD–RNA (Nudt12, Nudt16) or cleaving free NAD or NADH (Nudt13^*41*^, Nudt14^*39*^ and Nudt7^*42*^) (Figure S11). Potentially, some NudiXes may take over the function of the downregulated enzyme DXO and decap NAD–RNA. Surprisingly, Nudt12 was not expressed at all in either of the conditions (control and DXO KD). The expression levels of other enzymes were not significantly changed upon the knockdown of DXO. Therefore, we concluded that some unknown decapping enzyme should be activated and cleave NAD from RNA in DXO KD. On the other hand, overexpression of the decapping enzyme DXO led to a more than six–fold decrease in the level of NAD capping of sRNA (Figure 3F). Because we observed a decrease of NAD capping in U1 RNA from HIV– 1–infected cells in NAD captureSeq and LC–MS experiments and because U1 is important for HIV–1 infection, we were interested in knowing whether manipulated levels of DXO influence the NAD capping of this RNA. We observed a statistically non–significant decrease in NAD capping of U1 in MT4–DXO OE cells and an increase in MT4 DXO KD cells (Figure 3G, Figure S9D, Figure S12A–C). Together with the results on HIV–1 infectivity, these data suggest that overexpression of the DXO enzyme promotes HIV–1 replication through decreasing the level of NAD RNA capping. The result also indicates that the reduction of the NAD RNA cap increases HIV–1 production and that NAD–capped RNAs might play a role in antiviral defence.

In summary, we demonstrate that HIV–1 infection influences not only the total cellular NAD pool but also the NAD RNA capping of specific snRNAs (U1, U4ATAC, U5E and U7) and snoRNAs (SNORD3G, SNORD102, SNORA50A and SNORD3B). As U1 plays an important role in HIV–1 infection by binding to HIV–1 pre– mRNA, we investigated the capability of the NAD RNA cap to contribute to U1 RNA and HIV–1 pre–mRNA duplex formation. The presence of NAD on U1 leads to less stable RNA–RNA duplexes compared with TMG–capped U1. Additionally, we demonstrate that general NAD RNA capping is unfavorable for HIV–1 infection, as overexpression of the NAD decapping enzyme DXO results in decreased NAD capping and higher viral infectivity. Our work may provide an explanation for the finding that a polymorphism within the 5′ UTR region of the DXO gene in an African American cohort of HIV–1 patients^*22*^ is associated with an altered response to HIV–1 infection. Even though the mutation in the 5′ UTR region does not influence the protein sequence itself, it may influence the stability or translatability of RNA and thus the intracellular level of the DXO protein, which in turn alters the efficiency of HIV–1 replication. In conclusion, we hypothesize that NAD–capped sRNAs inhibit HIV–1 replication and that part of the virus’s strategy to overcome this inhibition is to reduce the pool of NAD–capped RNA. RNA modifications such as 6– methyladenosine^*43*^ and 2’–O–methylation^*44*^ of the HIV–1 genomic RNA were found to play an important role in HIV–1 infection. The NAD cap, thus, might be another crucial RNA modification. It is present on host RNAs, influenced by the infection and it has an impact on the viral infection itself. Moreover, we (and others) have also observed the depletion of total cellular NAD in other viral infections: Severe acute respiratory syndrome coronavirus 2 (SARS-CoV-2)^*45*^ and Herpes Simplex virus (HSV–1) (Figure S13) from different classes of viruses than retroviruses. It would be intriguing to study whether the change in total NAD results in a change of NAD capping on the same or similar RNAs as in the case of HIV–1, and whether the presence of the NAD cap negatively influences the viral infection. If it is a general mechanism, exploiting NAD–capped RNA for therapeutic purposes might present a future avenue of antiviral treatment.

## Acknowledgements

We are grateful to all members of the Cahova Group for their help and advice, particularly Z. Buchová for her help with the preparation of the DXO enzyme. We acknowledge funding from the Ministry of Education, Youth and Sports of the Czech Republic, programme ERC CZ (LL1603), the European Research Council Executive Agency (ERCEA) under the European Union’s Horizon Europe Framework Programme for Research and Innovation (grant agreement No. 101041374 – StressRNaction) and the Operational Programme Johannes Amos Comenius (OP JAC) project RNA for Therapy, reg. No. CZ.02.01.01/00/22_008/0004575 co–financed by the EU. D. S. and H.C. acknowledge support from the Czech Science Foundation (project No. 24–11157S). Computational resources were supplied by the project ‘e–Infrastruktura CZ’ (e–INFRA CZ LM2018140) supported by the Ministry of Education, Youth and Sports of the Czech Republic.

## Author contributions

B. B., A. Š., P. V. and H. C. designed the experiments and coordinated the project. J. P., J. W. and D. S. participated in the design of the experiments. B. B., J. F. P., A. Š., P. V., R. B., J. T., K. G., M.–B. M., L. B. and H. C. performed the experiments. K. G. and J. K. prepared TMG and coordinated the chemical synthesis. L. B. measured CD spectra. Z. K. and P. V. designed and coordinated the preparation of OE and KD DXO. L. G. performed all bioinformatic analyses. H. C. supervised the work. P. V., B. B. and H. C. wrote the paper.

## Supporting Information Available

This material is available free of charge via the Internet. Sequencing data are accessible in the GEO database under accession GSE191019.

