## Supplemental Information for "HIV–1 infection reduces NAD capping of host cell snRNA and snoRNA"

*Supplementary Table 1: Sequences of used oligonucleotides.*

| Name | **Sequence** (starting at 5´ end) |
|---|---|
| RNA 3’adaptor | rApp–CNNNNNNAGATCGGAAGAGCACACGTCTG–(C3) |
| Reverse transcription primer | CAGACGTGTGCTCTTCCGAT |
| cDNA anchor sense | p–CAGATCGGAAGAGCGTCGTGT–(C3) |
| cDNA anchor antisense | ACACGACGCTCTTCCGATCTGGG |
| NAD Seq Fwd PCR primer | AATGATACGGCGACCACCGAGATCTACACTCTTTCCCTACACGACGCTCTTCCGATCT |
| NAD Seq Rev PCR primer 1 | CAAGCAGAAGACGGCATACGAGATGCCTAAGTGACTGGAGTTCAGACGTGTGCTCTTCCG ATCT |
| NAD Seq Rev PCR primer 2 | CAAGCAGAAGACGGCATACGAGATTGGTCAGTGACTGGAGTTCAGACGTGTGCTCTTCCG ATCT |
| NAD Seq Rev PCR primer 3 | CAAGCAGAAGACGGCATACGAGATCACTGTGTGACTGGAGTTCAGACGTGTGCTCTTCCGA TCT |
| NAD Seq Rev PCR primer 4 | CAAGCAGAAGACGGCATACGAGATATTGGCGTGACTGGAGTTCAGACGTGTGCTCTTCCG ATCT |

| | |
|---|---|
| NAD Seq Rev PCR primer 5 | CAAGCAGAAGACGGCATACGAGATGATCTGGTGACTGGAGTTCAGACGTGTGCTCTTCCGATCT |
| NAD Seq Rev PCR primer 6 | CAAGCAGAAGACGGCATACGAGATTCAAGTGTGACTGGAGTTCAGACGTGTGCTCTTCCGATCT |
| NAD Seq Rev PCR primer 7 | CAAGCAGAAGACGGCATACGAGATCTGATCGTGACTGGAGTTCAGACGTGTGCTCTTCCGATCT |
| NAD Seq Rev PCR primer 8 | CAAGCAGAAGACGGCATACGAGATAAGCTAGTGACTGGAGTTCAGACGTGTGCTCTTCCGATCT |
| SNORD3G Fwd | GCGTTCTCTCCTGAGCATGA |
| SNORD3G Rev | ACCACTCAGACTGTGTCCTCT |
| SNORD3B Fwd | TCTGAACGTGTAGAGCACCG |
| SNORD3B Rev | CTCTCCCTCTCACTCCCCAA |
| SNORA50A Fwd | GCACTGCCTTTGAACCTGATG |
| SNORA50A Rev | GAGCAGTTCAGTTGAGAGCTG |
| SNORD102 Fwd | GACTGTTTTTTTTGATTGCTTG |
| SNORD102 Rev | AGCCGGTGAAATGTGT |
| U1 Fwd | TGGCAGGGGAGATACCATGA |
| U1 Rev | TACGCAGTCGAGTTTCCCAC |
| U4 ATAC Fwd | CCATCCTTTTCTTGGGGTTGC |
| U4 ATAC Rev | GCAAAAGCTCTAGTTGATGCGG |
| U5E Fwd | AGTTTCTCTTCATATCGCA |
| U5E Rev | AAAATGCAAGACCTCT |
| U7 Fwd | GTGTTACAGCTCTTTTAG |
| U7 Rev | TCCGGTAAAAAGCCAG |
| Fwd primer for U1 RNA template preparation | TAATACGACTCACTATTAGGCTTACCTGGCAGGGGAG |
| Rev primer for U1 RNA template preparation | CAGGGGAAAGCGCGAACGCAG |
| U1 RNA template 20 nt | TAATACGACTCACTATTAGGCTTACCTGGCAGGGGAG |
| HIV–1 mRNA 20 nt | TAATACGACTCACTATAGGGTAAGCCGTACATGTAAT |

| | |
|---|---|
| Biotinylated U1 probe | GCAATGGATAAGCCTCGCCC |
| DXO qPCR Fwd | CTTCTGCTCTGTGCTACGCA |
| DXO qPCR Rev | TGGCCAGGGCTGTGCATC |
| NUDT7 qPCR Fwd | CTAAGGCCCGCTTAAGAAAGTA |
| NUDT7 qPCR Rev | CGGACGGTGAACAACAAATG |
| NUDT12 qPCR Fwd | CTGGTGAAGTCCCGAGAGAG |
| NUDT12 qPCR Rev | TTGAGCTACAACCCCAGCTT |
| NUDT13 qPCR Fwd | ATCACGCTGGTGTCAGATGGGA |
| NUDT13 qPCR Rev | AACTTCTCGGCGGATGGTCTCT |
| NUDT14 qPCR Fwd | TTGGGAGGAGTGTGGCTACCAC |
| NUDT14 qPCR Rev | GGCATCTGTCACCTCTGTGTAG |
| NUDT16 qPCR Fwd | TACGCCATACTGATGCAGA |
| NUDT16 qPCR Rev | GTCCTCTAGGCTTCTGTCCT |
| ACTB Fwd | CACCAACTGGGACGACAT |
| ACTB Rev | ACAGCCTGGATAGCAACG |

*Supplementary Table 2: List of 94 sRNAs with enrichment in +ADPRC samples over –ADPRC control (corresponding to the step 3 in Figure S1B). Last three columns mark whether the sRNA passed (1) or did not pass (0) following filtering steps 4, 5 and 6 as described in Figure S1B.*

| shortRNA | Chr | Start | End | Strand | mean –HIV | mean +HIV | step4 | step5 | step6 |
|---|---|---|---|---|---|---|---|---|---|
| RNU5E–4P–201 | 1 | 11909808 | 11909927 | - | 1.06 | -0.57 | 1 | 1 | 1 |
| Gly | 1 | 16678271 | 16678341 | - | 1.09 | 1.27 | 0 | 0 | 0 |
| RNU6-40P-201 | 1 | 31497577 | 31497683 | - | 0.06 | 0.31 | 0 | 0 | 0 |
| SNORD3G-201 | 1 | 90657750 | 90657964 | + | 3.16 | 1.80 | 1 | 1 | 1 |
| SNORD21-201 | 1 | 92837289 | 92837383 | + | 0.66 | -0.20 | 1 | 1 | 0 |
| Cys | 1 | 93516277 | 93516349 | - | 1.08 | 0.46 | 0 | 0 | 0 |
| Gly | 1 | 121016845 | 121016915 | - | 1.50 | 1.21 | 0 | 0 | 0 |
| Asn | 1 | 145287766 | 145287839 | + | 0.31 | 1.01 | 0 | 0 | 0 |
| Gly | 1 | 146037061 | 146037132 | + | -0.08 | 1.54 | 0 | 0 | 0 |
| RNVU1-1-201 | 1 | 148362370 | 148362533 | - | 1.50 | 0.04 | 1 | 1 | 1 |
| RNVU1-30-201 | 1 | 149636766 | 149636929 | + | 0.81 | 0.21 | 1 | 0 | 0 |
| Val | 1 | 149708660 | 149708730 | - | 4.20 | 3.69 | 0 | 0 | 0 |
| Met | 1 | 153671250 | 153671321 | + | 1.25 | 1.32 | 0 | 0 | 0 |
| Ala | 2 | 27051214 | 27051286 | + | 0.36 | 1.20 | 1 | 1 | 1 |
| Ile | 2 | 42810536 | 42810628 | + | 1.09 | 0.61 | 0 | 0 | 0 |
| RNU4ATAC-201 | 2 | 121530881 | 121531007 | + | 1.80 | -0.53 | 1 | 1 | 1 |
| MIR425-201 | 3 | 49020148 | 49020234 | - | 0.78 | 0.22 | 0 | 0 | 0 |
| RNU2-64P-201 | 3 | 73110992 | 73111182 | + | 0.34 | 0.12 | 0 | 0 | 0 |
| Cys | 3 | 132231798 | 132231869 | - | 1.46 | 0.39 | 1 | 1 | 0 |
| Val | 3 | 169772230 | 169772302 | + | 1.20 | 0.45 | 0 | 0 | 0 |
| U3.44-201 | 4 | 158700691 | 158700909 | + | 2.37 | 1.45 | 0 | 0 | 0 |
| Leu | 5 | 181097474 | 181097555 | - | 0.72 | 1.09 | 0 | 0 | 0 |
| Val | 5 | 181188416 | 181188488 | - | 1.21 | 1.06 | 0 | 0 | 0 |
| Met | 6 | 26286526 | 26286597 | + | -0.07 | 1.28 | 1 | 1 | 0 |
| Val | 6 | 26538054 | 26538126 | + | 0.99 | 0.53 | 0 | 0 | 0 |
| Ile | 6 | 26554122 | 26554195 | + | 2.01 | 2.79 | 0 | 0 | 0 |
| Ala | 6 | 26571864 | 26571936 | - | 0.94 | 0.10 | 1 | 1 | 0 |
| Tyr | 6 | 26575570 | 26575659 | + | 1.02 | 0.32 | 1 | 1 | 0 |
| Ile | 6 | 26720992 | 26721065 | - | 0.26 | -0.21 | 1 | 1 | 0 |
| Ile | 6 | 27237571 | 27237644 | - | 0.95 | -0.03 | 0 | 0 | 0 |
| Ser | 6 | 27495814 | 27495895 | + | 0.60 | 0.82 | 0 | 0 | 0 |
| Ser | 6 | 27503039 | 27503120 | + | 0.80 | 1.46 | 0 | 0 | 0 |
| Lys | 6 | 27591814 | 27591886 | - | 1.95 | 1.50 | 0 | 0 | 0 |
| Met | 6 | 27592821 | 27592892 | - | -0.33 | 0.52 | 1 | 1 | 0 |
| Arg | 6 | 27670565 | 27670637 | - | 0.78 | 0.68 | 0 | 0 | 0 |
| Gln | 6 | 27791356 | 27791427 | - | 0.48 | 0.99 | 0 | 0 | 0 |
| Gln | 6 | 27795861 | 27795932 | - | 1.75 | 1.73 | 0 | 0 | 0 |

| | | | | | | | | | |
|---|---|---|---|---|---|---|---|---|---|
| Met | 6 | 27902493 | 27902564 | - | 0.51 | 0.46 | 1 | 1 | 0 |
| Ala | 6 | 28607156 | 28607227 | + | 0.95 | 0.81 | 0 | 0 | 0 |
| Ala | 6 | 28710589 | 28710660 | + | 0.81 | 0.79 | 0 | 0 | 0 |
| Ala | 6 | 28758364 | 28758435 | - | 3.66 | 3.53 | 0 | 0 | 0 |
| Ala | 6 | 28789770 | 28789841 | - | 0.59 | 0.36 | 0 | 0 | 0 |
| Ala | 6 | 28802800 | 28802870 | - | 0.86 | 0.21 | 1 | 1 | 0 |
| Ala | 6 | 28817235 | 28817306 | - | 1.26 | 0.49 | 0 | 0 | 0 |
| Ala | 6 | 28838444 | 28838515 | - | -0.23 | 0.89 | 1 | 0 | 0 |
| Leu | 6 | 28943622 | 28943703 | - | 0.06 | 0.91 | 1 | 1 | 0 |
| Met | 6 | 28944575 | 28944647 | + | 0.56 | 0.84 | 0 | 0 | 0 |
| Lys | 6 | 28951029 | 28951101 | + | -0.17 | 1.11 | 1 | 1 | 0 |
| Leu | 6 | 28989002 | 28989083 | + | -0.48 | 1.01 | 1 | 1 | 0 |
| Cys | 7 | 149595214 | 149595285 | - | -0.23 | 0.79 | 1 | 0 | 0 |
| Y_RNA.402-201 | 8 | 99527826 | 99527927 | + | -0.40 | 0.72 | 1 | 0 | 0 |
| RNU1-35P-201 | 8 | 135742343 | 135742495 | - | 1.56 | 0.31 | 1 | 1 | 1 |
| Asn | 10 | 22229509 | 22229582 | - | 0.56 | 1.86 | 0 | 0 | 0 |
| Ser | 10 | 67764503 | 67764584 | + | -0.79 | 1.13 | 1 | 1 | 0 |
| Y_RNA.234-201 | 11 | 3663815 | 3663911 | - | 0.35 | 1.45 | 1 | 1 | 1 |
| Lys | 11 | 59556429 | 59556501 | + | 0.16 | 0.77 | 1 | 1 | 0 |
| Lys | 11 | 59560335 | 59560407 | - | 1.04 | 0.54 | 0 | 0 | 0 |
| RNU7-1-201 | 12 | 6943816 | 6943878 | + | 1.14 | 0.29 | 1 | 1 | 1 |
| Trp | 12 | 98504252 | 98504323 | + | -0.08 | 1.81 | 1 | 1 | 1 |
| Ala | 12 | 124921755 | 124921826 | - | 1.95 | 1.85 | 0 | 0 | 0 |
| SNORA49-201 | 12 | 132031224 | 132031359 | + | -0.13 | 1.12 | 1 | 1 | 0 |
| SNORD102-201 | 13 | 27255064 | 27255135 | + | 0.82 | -0.52 | 1 | 1 | 1 |
| Asn | 13 | 30673964 | 30674037 | - | 0.96 | 1.38 | 1 | 1 | 1 |
| Glu | 13 | 41060738 | 41060809 | - | 1.66 | 0.96 | 0 | 0 | 0 |
| MIR19B1-201 | 13 | 91351192 | 91351278 | + | -0.55 | 0.74 | 1 | 1 | 1 |
| Phe | 13 | 94549650 | 94549722 | - | 1.02 | 0.79 | 0 | 0 | 0 |
| Pro | 14 | 20609336 | 20609407 | - | 0.93 | 0.90 | 0 | 0 | 0 |
| Pro | 14 | 20684016 | 20684087 | + | 1.42 | 0.06 | 0 | 0 | 0 |
| Lys | 14 | 58239895 | 58239967 | - | 1.26 | 0.87 | 0 | 0 | 0 |
| Y_RNA.351-201 | 14 | 93397139 | 93397234 | - | 0.30 | 1.08 | 0 | 0 | 0 |
| Ile | 14 | 102317092 | 102317165 | + | 0.23 | 1.18 | 1 | 1 | 0 |
| Leu | 16 | 22297140 | 22297221 | + | 1.33 | 0.33 | 0 | 0 | 0 |
| Leu | 16 | 57299951 | 57300033 | + | 1.09 | 1.23 | 0 | 0 | 0 |
| SNORA50A-201 | 16 | 58559796 | 58559929 | - | 0.52 | -0.71 | 1 | 1 | 1 |
| Y_RNA.793-201 | 16 | 66550457 | 66550567 | + | 0.04 | 1.41 | 1 | 0 | 0 |
| Lys | 16 | 73478317 | 73478389 | - | 0.88 | 0.23 | 0 | 0 | 0 |
| Gln | 17 | 8119752 | 8119823 | + | 1.59 | 1.36 | 0 | 0 | 0 |
| Gly | 17 | 8221548 | 8221619 | + | 1.59 | 1.49 | 0 | 0 | 0 |

| | | | | | | | | | |
|---|---|---|---|---|---|---|---|---|---|
| Ile | 17 | 8226991 | 8227064 | - | 1.13 | 0.56 | 0 | 0 | 0 |
| SNORD3B-2-201 | 17 | 19063346 | 19064136 | - | 4.04 | 0.21 | 1 | 1 | 1 |
| SNORD3C-201 | 17 | 19189665 | 19190245 | - | 0.42 | 0.82 | 0 | 0 | 0 |
| Thr | 17 | 31550074 | 31550145 | + | 0.06 | 1.01 | 1 | 0 | 0 |
| Cys | 17 | 38861684 | 38861755 | - | 1.08 | 0.39 | 1 | 1 | 1 |
| RNU1-42P-201 | 17 | 48949361 | 48949520 | - | 1.90 | 0.32 | 1 | 1 | 1 |
| U3.18-201 | 17 | 58631641 | 58631836 | - | 2.52 | 1.03 | 0 | 0 | 0 |
| AC025362.1-201 | 17 | 64146337 | 64146471 | + | 0.40 | 0.50 | 0 | 0 | 0 |
| SNORA50C-201 | 17 | 64146339 | 64146471 | + | 0.40 | 0.50 | 0 | 0 | 0 |
| Met | 17 | 82494721 | 82494792 | - | 1.33 | 2.35 | 0 | 0 | 0 |
| RNU6-346P-201 | 18 | 76800664 | 76800774 | - | -0.03 | -0.10 | 0 | 0 | 0 |
| RNU6-2-201 | 19 | 1021522 | 1021628 | + | 1.59 | -0.44 | 1 | 1 | 0 |
| Pseudo | 19 | 41242237 | 41242309 | - | 0.36 | 1.16 | 0 | 0 | 0 |
| SNORD88A-201 | 19 | 50799442 | 50799532 | - | -0.15 | 0.80 | 1 | 1 | 0 |
| SNORD86-201 | 20 | 2656097 | 2656182 | + | 0.56 | 0.78 | 0 | 0 | 0 |
| RNU6-759P-201 | 20 | 35647328 | 35647430 | - | -0.52 | 0.57 | 1 | 0 | 0 |

*Supplementary Table 3: RNA-seq bioinformatic analysis of sRNAs in noninfected and HIV-1 infected cells.*

| RNA | Chromosome | Start | End | Probe Strand | P-value (DESeq stats p<0.05 after correction) | FDR (DESeq stats p<0.05 after correction) | Log2 Fold Change (DESeq stats p<0.05 after correction) | Shrunk Log2 Fold Change (DESeq stats p<0.05 after correction) | noninfected | infected |
|---|---|---|---|---|---|---|---|---|---|---|
| Lys | 6 | 27576067 | 27576139 | + | 2.35E-09 | 5.05E-07 | 2.816879 | 2.297963 | 5.493564 | 3.064846 |
| Y_RNA.392 | 12 | 17710934 | 17711046 | + | 1.53E-04 | 0.006597 | 3.753169 | 1.921711 | 3.249257 | 0.166602 |
| Glu | 13 | 41060738 | 41060809 | - | 3.78E-06 | 2.80E-04 | 2.506555 | 1.920643 | 5.962084 | 3.714684 |
| Y_RNA.562 | X | 96701507 | 96701619 | - | 3.76E-06 | 2.80E-04 | 2.316154 | 1.840333 | 4.834023 | 2.816057 |
| Y_RNA.83 | 11 | 93719603 | 93719715 | + | 1.61E-04 | 0.006656 | 2.307293 | 1.66392 | 4.315581 | 2.288709 |
| Val | 6 | 26538054 | 26538126 | + | 2.46E-04 | 0.009051 | 2.19048 | 1.596283 | 6.385199 | 4.111965 |
| Pro | 14 | 20633006 | 20633077 | + | 4.04E-07 | 4.57E-05 | 1.769528 | 1.571224 | 9.919452 | 8.327454 |
| Lys | 17 | 8119155 | 8119227 | + | 8.60E-07 | 8.40E-05 | 1.764935 | 1.557631 | 8.722684 | 7.186026 |
| Asn | 17 | 38751781 | 38751854 | - | 2.80E-05 | 0.001585 | 1.851575 | 1.540986 | 6.109306 | 4.159446 |
| Y_RNA.17 | 6 | 99642237 | 99642347 | - | 3.89E-04 | 0.012873 | 2.063833 | 1.533198 | 3.940254 | 2.12569 |
| Gly | 2 | 1.56E+08 | 1.56E+08 | - | 2.63E-05 | 0.001534 | 1.771341 | 1.496111 | 8.240386 | 6.624817 |
| Y_RNA.573 | 10 | 1.25E+08 | 1.25E+08 | - | 7.16E-05 | 0.003273 | 1.801234 | 1.484377 | 6.81489 | 5.317549 |
| Pro | 16 | 3158922 | 3158993 | + | 2.58E-05 | 0.001534 | 1.717011 | 1.464559 | 8.497743 | 6.896557 |
| Y_RNA.518 | 20 | 18113467 | 18113579 | - | 4.92E-05 | 0.002349 | 1.726967 | 1.454194 | 7.173159 | 5.70289 |
| Y_RNA.516 | 10 | 36626015 | 36626138 | - | 5.81E-06 | 4.03E-04 | 1.639518 | 1.443974 | 8.316263 | 6.900973 |
| Y_RNA.10 | 1 | 2.41E+08 | 2.41E+08 | + | 1.74E-04 | 0.006938 | 1.752024 | 1.429235 | 5.421762 | 3.944697 |
| MIR4493 | 11 | 1.23E+08 | 1.23E+08 | - | 0.001641 | 0.043529 | 2.047502 | 1.42916 | 4.573041 | 2.746447 |

| | | | | | | | | | | |
|---|---|---|---|---|---|---|---|---|---|---|
| MIR29B1 | 7 | 1.31E+08 | 1.31E+08 | - | 1.75E-06 | 1.51E-04 | 1.585097 | 1.423341 | 8.343235 | 6.928911 |
| Y_RNA.555 | 5 | 1.09E+08 | 1.09E+08 | + | 3.75E-04 | 0.012691 | 1.79408 | 1.41903 | 5.881616 | 4.369372 |
| Y_RNA.476 | 4 | 37699895 | 37700002 | - | 4.74E-04 | 0.015445 | 1.816481 | 1.418557 | 5.325714 | 3.759005 |
| Y_RNA.32 | 1 | 85435175 | 85435284 | - | 2.35E-04 | 0.009003 | 1.752486 | 1.418348 | 5.715662 | 4.190845 |
| Gly | 1 | 1.62E+08 | 1.62E+08 | - | 3.04E-04 | 0.010703 | 1.69115 | 1.377953 | 8.010199 | 6.396339 |
| Y_RNA.435 | 14 | 63621760 | 63621871 | + | 3.49E-05 | 0.001875 | 1.517128 | 1.331911 | 10.82305 | 9.56219 |
| Pro | 14 | 20609336 | 20609407 | - | 9.84E-04 | 0.028186 | 1.620699 | 1.295579 | 6.612308 | 5.13773 |
| Y_RNA.70 | 1 | 28881726 | 28881835 | - | 3.11E-04 | 0.010786 | 1.514026 | 1.280204 | 8.139081 | 6.858013 |
| Y_RNA.696 | 17 | 60748940 | 60749046 | - | 5.79E-04 | 0.018457 | 1.547087 | 1.279207 | 5.337646 | 3.977821 |
| Pro | 14 | 20684016 | 20684087 | + | 8.17E-05 | 0.003657 | 1.416925 | 1.249693 | 8.480556 | 7.334147 |
| MIR1468 | X | 63786002 | 63786087 | - | 0.001475 | 0.039635 | 1.521453 | 1.229377 | 6.145439 | 4.852688 |
| Ala | 12 | 1.25E+08 | 1.25E+08 | - | 0.001881 | 0.047545 | 1.514852 | 1.216813 | 5.429174 | 4.101664 |
| Trp | 17 | 19508181 | 19508252 | + | 6.03E-04 | 0.018497 | 1.413615 | 1.202712 | 5.418914 | 4.264954 |
| U3.18 | 17 | 58631641 | 58631836 | - | 5.92E-04 | 0.018457 | 1.353132 | 1.16649 | 4.399987 | 3.071916 |
| Y_RNA.653 | 9 | 70311607 | 70311701 | - | 1.93E-05 | 0.001221 | 1.0845 | 1.016812 | 7.039675 | 6.11551 |
| MIR25 | 7 | 1E+08 | 1E+08 | - | 1.70E-05 | 0.001108 | 1.074476 | 1.009354 | 17.40652 | 16.49354 |
| MIR942 | 1 | 1.17E+08 | 1.17E+08 | + | 0.00207 | 0.049986 | 0.885955 | 0.816154 | 9.41885 | 8.683841 |
| MIR196A2 | 12 | 53991738 | 53991847 | + | 0.001445 | 0.039304 | 0.844595 | 0.787354 | 7.89752 | 7.222378 |
| MIR16-2 | 3 | 1.6E+08 | 1.6E+08 | + | 2.42E-04 | 0.009051 | 0.820448 | 0.780155 | 14.21309 | 13.55952 |
| MIR483 | 11 | 2134134 | 2134209 | - | 2.53E-04 | 0.009051 | 0.764608 | 0.731597 | 12.90623 | 12.35329 |
| SNORD57 | 20 | 2656939 | 2657010 | + | 0.001235 | 0.034462Z^Y | -0.60285 | -0.58191 | 12.73658 | 13.57901 |

| | | | | | | | | | | |
|---|---|---|---|---|---|---|---|---|---|---|
| SNORD83B | 22 | 39313819 | 39313911 | - | 7.78E-04 | 0.023228 | -0.83373 | -0.78387 | 9.227509 | 10.25942 |
| SNORD13 | 8 | 33513475 | 33513578 | + | 0.00186 | 0.047545 | -0.93499 | -0.85516 | 10.00742 | 11.0756 |
| SNORD45A | 1 | 75787889 | 75787972 | + | 9.82E-04 | 0.028186 | -1.03765 | -0.94119 | 9.279798 | 10.50702 |
| Gly | 21 | 17454789 | 17454859 | - | 3.59E-05 | 0.001884 | -1.02155 | -0.96085 | 13.51185 | 14.69965 |
| Vault.4 | 5 | 1.36E+08 | 1.36E+08 | - | 7.14E-07 | 7.30E-05 | -1.0333 | -0.98889 | 7.70246 | 9.068642 |
| RNU12 | 22 | 42615244 | 42615393 | + | 0.002065 | 0.049986 | -1.26534 | -1.07728 | 9.171878 | 10.57052 |
| SNORD89 | 2 | 1.01E+08 | 1.01E+08 | - | 0.001976 | 0.04881 | -1.27784 | -1.08617 | 8.719049 | 10.17037 |
| SNORA75 | 2 | 2.31E+08 | 2.31E+08 | - | 0.001431 | 0.039304 | -1.28305 | -1.09899 | 7.604465 | 9.045101 |
| SNORD94 | 2 | 86135870 | 86136006 | + | 4.66E-05 | 0.002276 | -1.23008 | -1.12428 | 5.972054 | 7.395618 |
| SNORD111 | 16 | 70538005 | 70538098 | + | 2.40E-07 | 2.86E-05 | -1.19338 | -1.13099 | 9.531701 | 11.17057 |
| SNORD67 | 11 | 46762389 | 46762499 | - | 0.001821 | 0.047545 | -1.38906 | -1.15314 | 5.303498 | 7.050073 |
| SNORA48 | 17 | 7574713 | 7574847 | + | 7.18E-04 | 0.021739 | -1.35028 | -1.16061 | 3.398652 | 4.979776 |
| Y_RNA.48 | 15 | 74983274 | 74983376 | - | 1.66E-04 | 0.006727 | -1.31823 | -1.17055 | 5.501172 | 6.975471 |
| SNORD116-23 | 15 | 25091786 | 25091877 | + | 1.29E-04 | 0.005649 | -1.34045 | -1.18943 | 5.640678 | 7.250891 |
| Lys | 16 | 3191501 | 3191573 | + | 3.78E-04 | 0.012691 | -1.51094 | -1.27303 | 6.955541 | 8.568355 |
| Lys | 14 | 58239895 | 58239967 | - | 1.25E-06 | 1.14E-04 | -1.38407 | -1.27649 | 10.25814 | 11.86491 |
| VTRNA3-1P | X | 53462209 | 53462310 | + | 2.64E-05 | 0.001534 | -1.47651 | -1.30964 | 4.843836 | 6.507765 |
| SNORD21 | 1 | 92837289 | 92837383 | + | 7.98E-04 | 0.023501 | -1.64168 | -1.31465 | 12.66439 | 14.27262 |
| SNORD92 | 2 | 28913664 | 28913748 | + | 5.27E-05 | 0.002463 | -1.53607 | -1.33652 | 8.04824 | 9.862964 |
| Met | 8 | 1.23E+08 | 1.23E+08 | - | 0.001838 | 0.047545 | -1.96982 | -1.39201 | 3.748004 | 5.573608 |
| SNORD3A | 17 | 19188016 | 19188714 | + | 0.001028 | 0.029057 | -2.1672 | -1.48575 | 4.595406 | 6.821272 |

| | | | | | | | | | | |
|---|---|---|---|---|---|---|---|---|---|---|
| Lys | 18 | 46089305 | 46089377 | - | 0.001934 | 0.048336 | -2.52625 | -1.4992 | 1.961316 | 4.266753 |
| VTRNA1-1 | 5 | 1.41E+08 | 1.41E+08 | + | 3.07E-11 | 1.29E-08 | -1.58815 | -1.49956 | 10.43433 | 12.28405 |
| SNORD15A | 11 | 75400391 | 75400538 | + | 1.44E-05 | 9.70E-04 | -1.78016 | -1.51552 | 13.30564 | 15.23064 |
| Glu | 15 | 26082234 | 26082305 | - | 1.57E-04 | 0.006605 | -1.97173 | -1.53816 | 5.604237 | 7.528315 |
| Y_RNA.266 | 10 | 88585638 | 88585733 | - | 3.19E-05 | 0.001755 | -1.94672 | -1.5885 | 4.233105 | 6.390816 |
| MIR1246 | 2 | 1.77E+08 | 1.77E+08 | - | 4.45E-05 | 0.002226 | -2.04627 | -1.62322 | 6.862081 | 9.065045 |
| RNU4-2 | 12 | 1.2E+08 | 1.2E+08 | - | 3.71E-06 | 2.80E-04 | -1.92791 | -1.63456 | 5.547839 | 7.616869 |
| RNU2-29P | 7 | 53776136 | 53776324 | + | 2.51E-04 | 0.009051 | -2.29145 | -1.63794 | 1.858093 | 4.250437 |
| SNORD14C | 11 | 1.23E+08 | 1.23E+08 | - | 2.16E-06 | 1.78E-04 | -1.92397 | -1.64357 | 6.933432 | 9.346354 |
| Y_RNA.297 | 14 | 51253933 | 51254023 | - | 2.09E-04 | 0.008166 | -2.55949 | -1.70104 | 5.781422 | 8.875075 |
| SNORD14D | 11 | 1.23E+08 | 1.23E+08 | - | 3.61E-11 | 1.29E-08 | -1.85959 | -1.71942 | 8.261262 | 10.32275 |
| SNORA13 | 5 | 1.12E+08 | 1.12E+08 | + | 1.09E-08 | 1.80E-06 | -1.95549 | -1.74473 | 6.068072 | 8.238724 |
| MIR122 | 18 | 58451074 | 58451158 | + | 3.78E-05 | 0.001933 | -2.50641 | -1.80668 | 5.206521 | 7.621225 |
| MIR215 | 1 | 2.2E+08 | 2.2E+08 | - | 1.50E-08 | 2.19E-06 | -2.06559 | -1.81535 | 4.276061 | 6.475221 |
| SNORD14A | 11 | 17074654 | 17074744 | - | 4.30E-06 | 3.08E-04 | -2.28803 | -1.82555 | 3.581273 | 6.048557 |
| RN7SKP79 | 5 | 6848127 | 6848462 | - | 5.93E-04 | 0.018457 | -3.80537 | -1.85681 | -2.12032 | 1.281619 |
| SNORD97 | 11 | 10801467 | 10801608 | - | 4.66E-12 | 2.50E-09 | -2.20139 | -1.99241 | 8.977421 | 11.27018 |
| Glu | 13 | 44917927 | 44917998 | - | 5.99E-07 | 6.43E-05 | -2.54868 | -2.00304 | 8.210685 | 10.91435 |
| RNU2-3P | 15 | 95745804 | 95745994 | + | 1.27E-06 | 1.14E-04 | -2.6645 | -2.02291 | 6.398067 | 9.188661 |
| MIR3609 | 7 | 98881650 | 98881729 | + | 1.81E-07 | 2.29E-05 | -2.56659 | -2.05033 | 5.991253 | 8.65035 |
| SNORD14E | 11 | 1.23E+08 | 1.23E+08 | - | 3.23E-09 | 6.30E-07 | -2.40956 | -2.05716 | 5.503731 | 8.25771 |

| | | | | | | | | | | |
|---|---|---|---|---|---|---|---|---|---|---|
| Telomerase-vert.1 | 3 | 1.7E+08 | 1.7E+08 | - | 1.24E-07 | 1.66E-05 | -2.71506 | -2.13033 | 3.144856 | 5.92663 |
| SNORD7 | 17 | 35573657 | 35573753 | + | 1.53E-08 | 2.19E-06 | -2.80182 | -2.23003 | 7.432362 | 10.33509 |
| U8.4 | 11 | 1.23E+08 | 1.23E+08 | + | 1.14E-09 | 2.72E-07 | -2.69111 | -2.23639 | 6.508183 | 9.262581 |
| RNY4P9 | 13 | 49908634 | 49908730 | - | 4.73E-10 | 1.27E-07 | -2.66775 | -2.24243 | 5.05719 | 7.913553 |
| Y_RNA.255 | 9 | 1.33E+08 | 1.33E+08 | - | 4.55E-09 | 8.15E-07 | -3.32347 | -2.48621 | 4.60583 | 8.056031 |
| SCARNA12 | 12 | 6967337 | 6967606 | - | 4.36E-18 | 4.68E-15 | -3.71242 | -3.11486 | 5.573442 | 9.542391 |
| SNORD3C | 17 | 19189665 | 19190245 | - | 2.41E-19 | 5.18E-16 | -4.54611 | -3.58749 | 1.042004 | 6.376476 |
| RN7SL397P | 3 | 1.2E+08 | 1.2E+08 | + | 5.52E-11 | 1.70E-08 | -8.70295 | -3.76366 | -3.10515 | 4.363092 |
| SNORD3D | 17 | 19112419 | 19112636 | - | 3.17E-13 | 2.27E-10 | -8.2338 | -4.75268 | -1.40851 | 7.710541 |

### Supplementary figures:

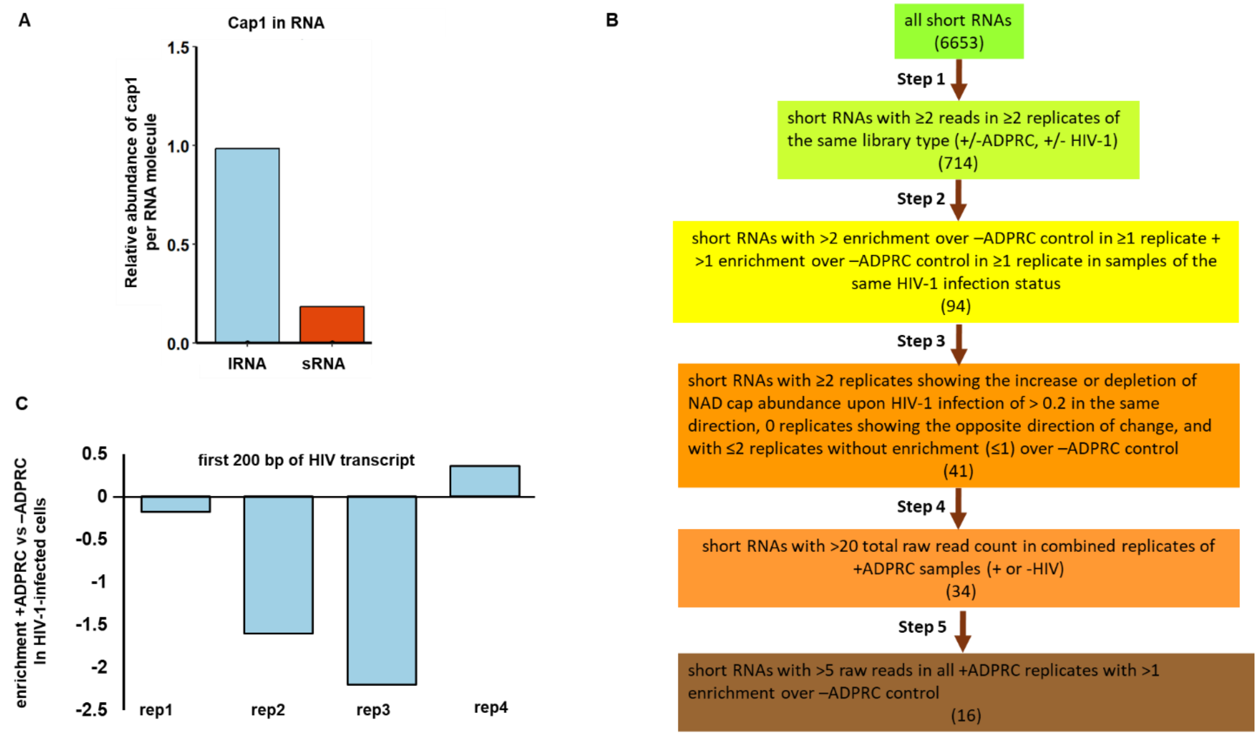


*Figure S 1: **Preparation for NAD captureSeq and NAD captureSeq analysis.** (A) LC-MS analysis of lRNA and sRNA from control cells digested by Nuclease P1. The intensity of signal of cap1 (m$^7$Gp$_3$Am) was normalized per amount of digested RNA (50 µg) and average length of sRNA (100 nt) and lRNA (2000 nt). (B) Bioinformatical analysis workflow describing individual filtering steps with the number of sRNA candidates after each step. (C) The enrichment (+ADPRC vs -ADPRC) of HIV–1 transcripts.*

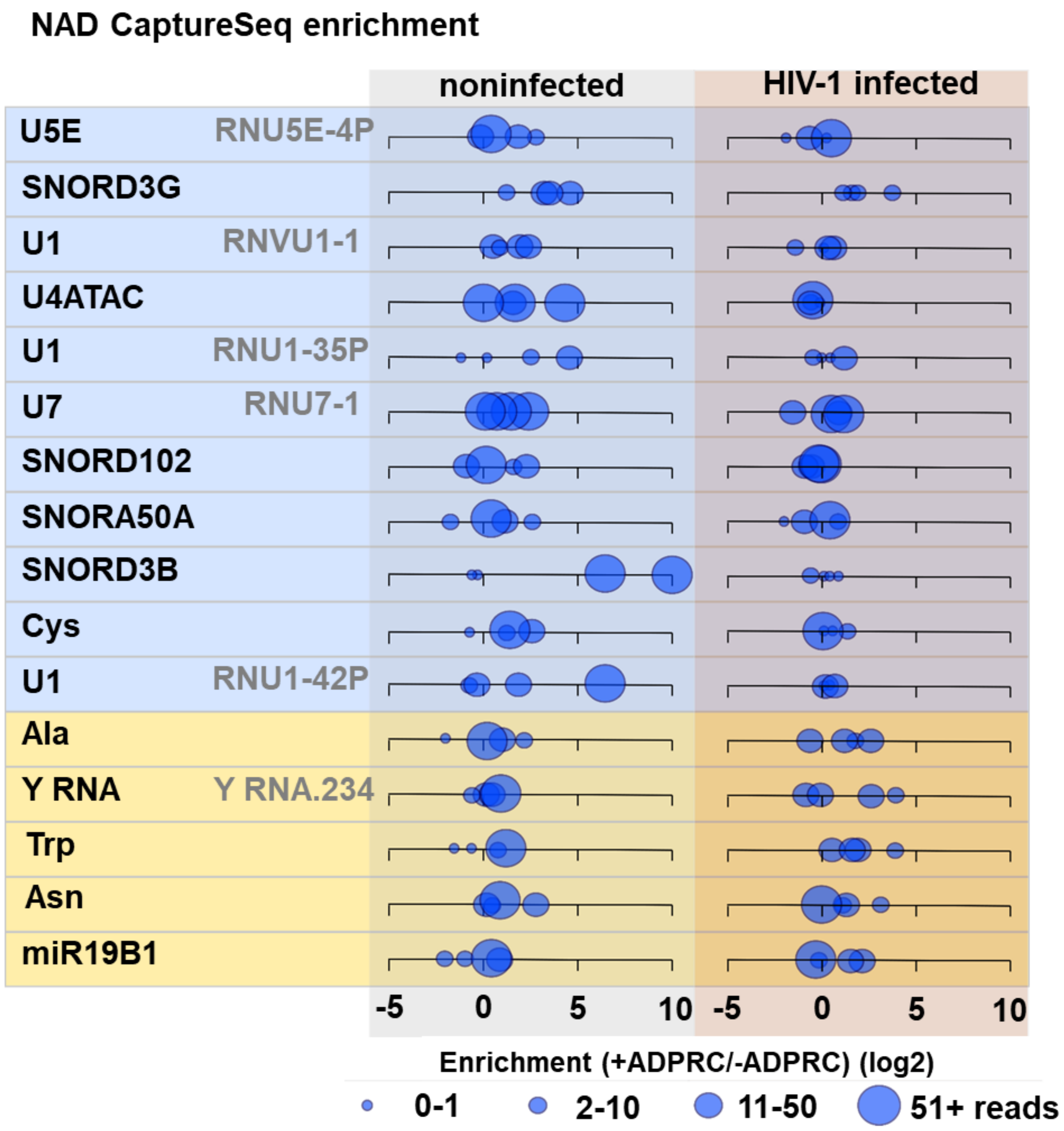


*Figure S 2:* ***Enrichment of RNAs in NAD captureSeq analysis.*** *The blue panel represents the RNAs enriched in control cells, whereas the yellow panel represents RNAs enriched in HIV–1 infected cells.*

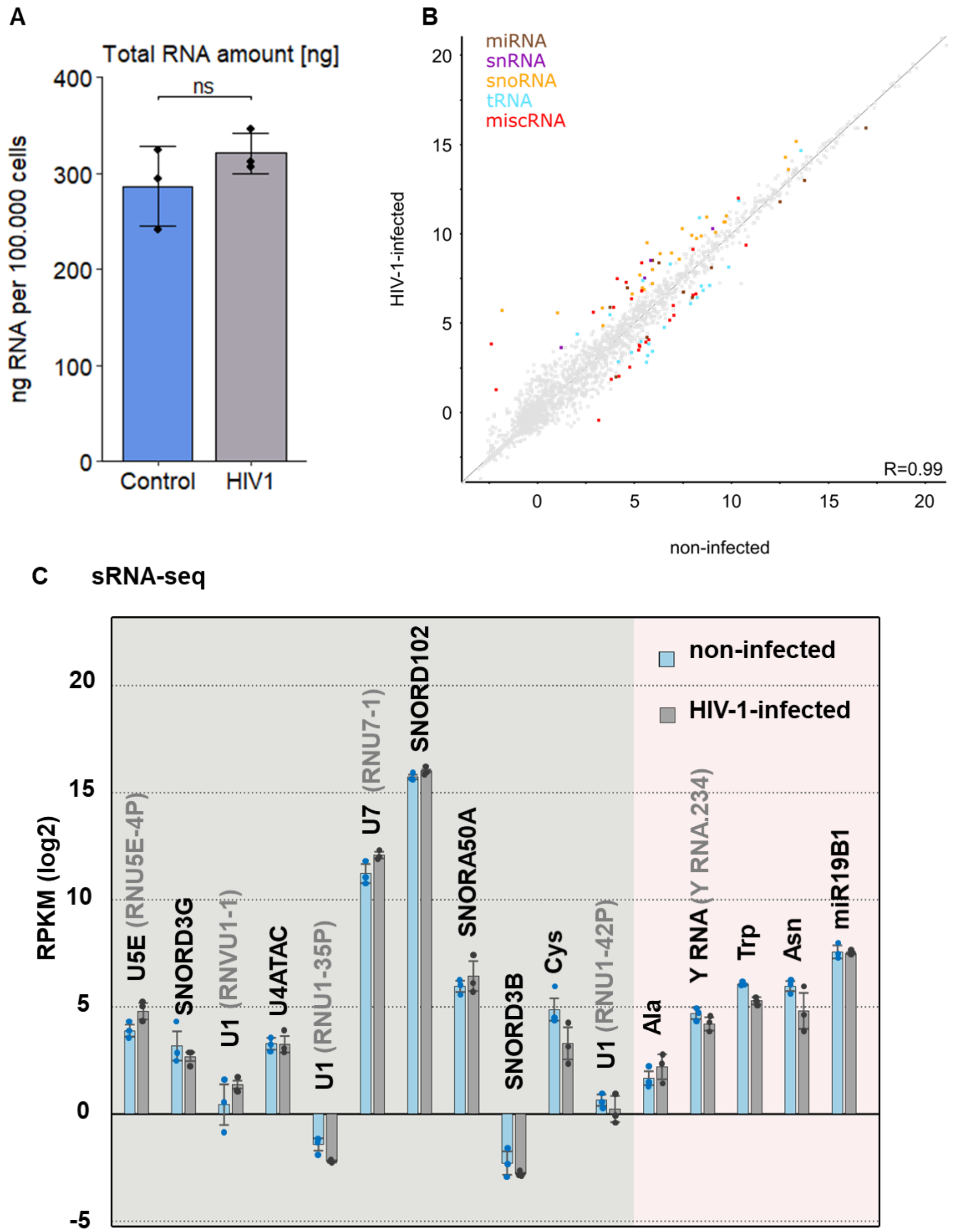


*Figure S 3: **Observed changes in NAD capping are not caused by different of amount of RNA or differential expression.** (A) Amount of total RNA isolated from control and HIV–1 infected. The error bar represents the standard deviation of the mean and black dots individual measurement from biological triplicate. B) All sRNAs analysis (miRNA, miscRNA, tRNA, snRNA, snoRNA, rRNA). Total sRNAs: 7624; Expressed (in sRNA datasets): 2014 (cut–off 0.5 in at least one condition); Significantly changing noninfected vs infected (in sRNA datasets): 89 (upregulated after infection: 52, downregulated after infection: 37) – highlighted by colour coding in the scatter plot. In general, sRNAs expression does not appear to be substantially altered after HIV infection. In scatter plot, dots below and to the right from the line are more expressed in non–infected, above and to the left more in HIV–1–infected MT–4 cells. snRNAs and snoRNAs that we identified as having decreased NAD cap after HIV infection do not have decreased expression after infection (no snRNAs and snoRNAs have decreased expression following HIV–1 infection). (C) sRNA–seq analysis of identified candidate sRNAs in control and HIV–1–infected cells (prepared in biological triplicates).*

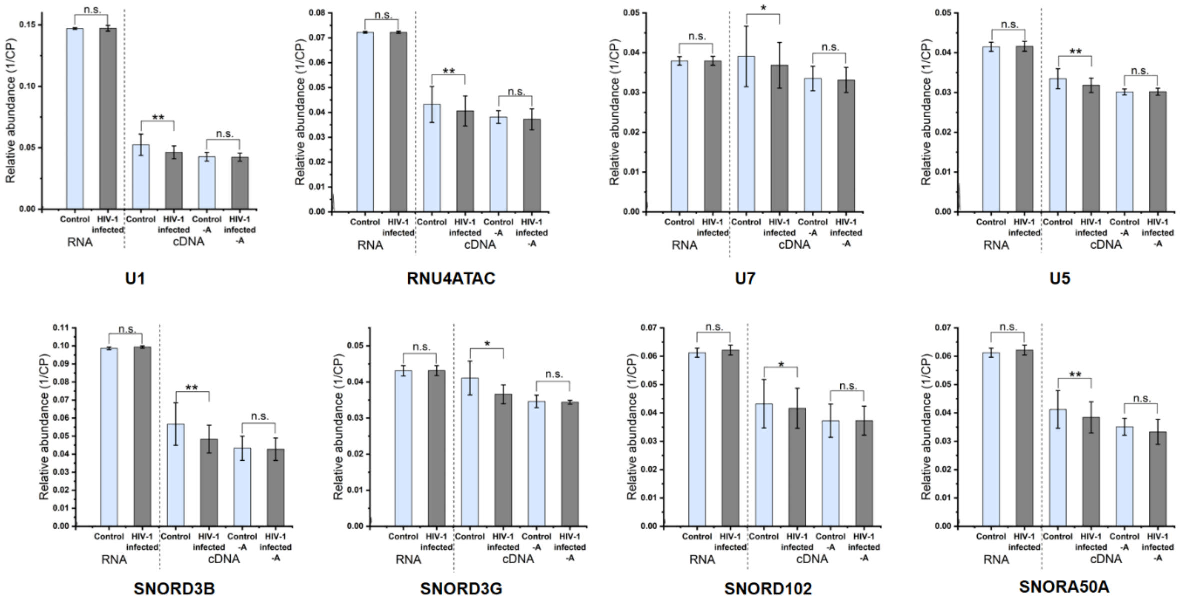


*Figure S 4:* ***RT–qPCR of isolated RNA and cDNA from NAD captureSeq library.*** *Measurement of relative abundance of each candidate sRNA in control and HIV–1 infected cells by RT–PCR and relative abundance of the corresponding cDNA in samples after NAD captureSeq protocol of control and HIV–1 infected cells.*

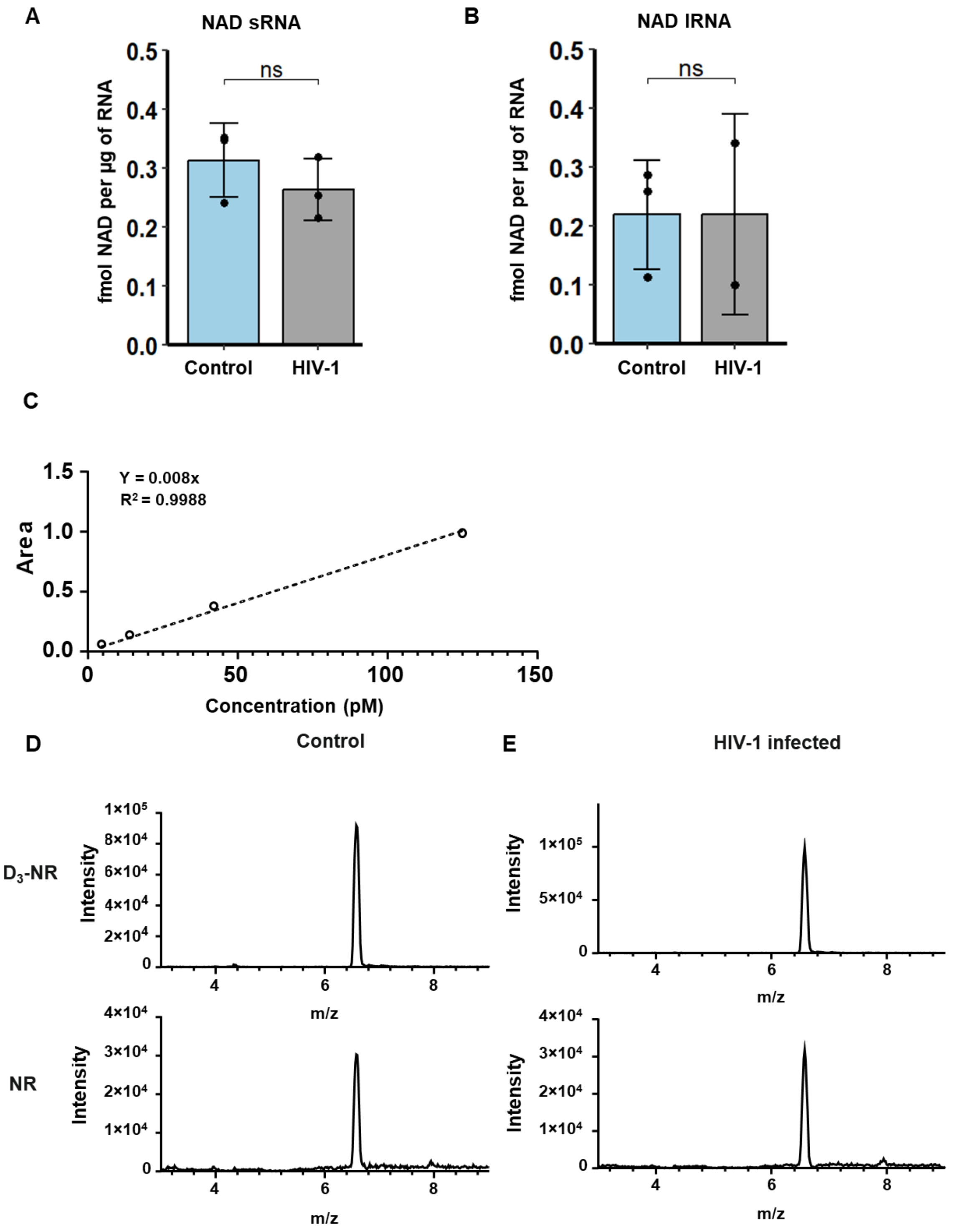


*Figure S 5:* ***LC–MS analysis of isolated sRNA and lRNA from control and HIV–1 infected cells.*** *(A) Absolute quantification of NAD RNA cap in fraction of sRNA from control and HIV–1 infected cells. (B) Absolute quantification of NAD RNA cap in fraction of lRNA from control and HIV–1 infected cells. (C) External calibration curve of NR analyte created from 4.6, 14, 42 and 125 pM concentraction. Each point is normalized to $D_3$–NR internal standard. (D) Extracted ion chromatograms of Tof–MRM transitions 258 →126 for $D_3$–NR used as internal standard and 255→123 for analysed NR in RNA from control cells. (E) Extracted ion chromatograms of Tof–MRM transitions 258 →126 for $D_3$–NR used as internal standard and 255→123 for analysed NR in RNA from HIV–1 infected cells.*

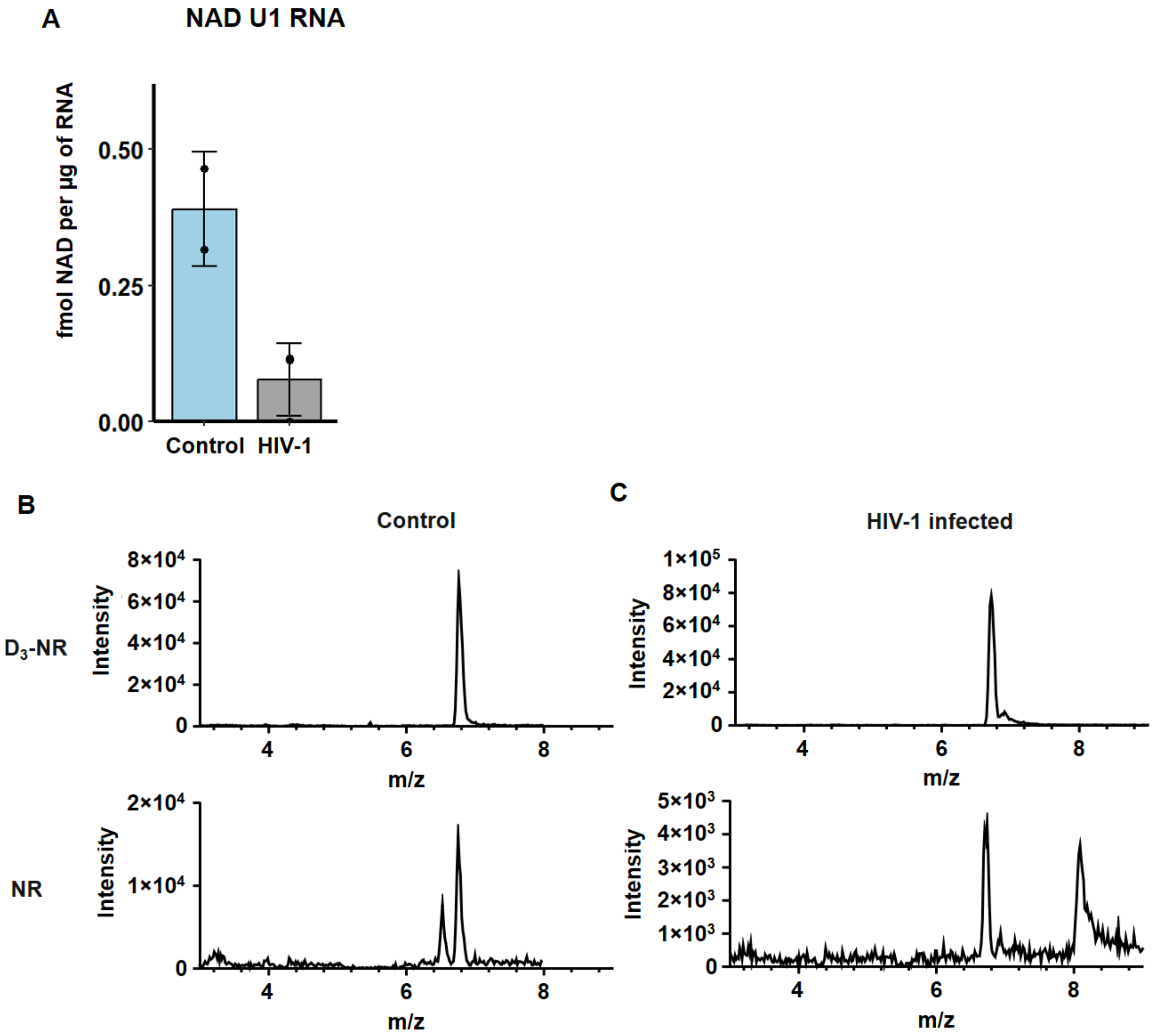


*Figure S 6:* ***LC–MS analysis of isolated U1 RNA pulled down from control and HIV–1 infected cells.*** *(A) Absolute quantification of NAD RNA cap in U1 pulled down from control and HIV–1 infected cells. (B) Extracted ion chromatograms of Tof–MRM transitions 258 →126 for $D_3$–NR used as internal standard and 255→123 for analysed NR in U1 RNA from control cells. (C) Extracted ion chromatograms of Tof–MRM transitions 258 →126 for $D_3$–NR used as internal standard and 255→123 for analysed NR in U1 RNA from HIV–1 infected cells.*

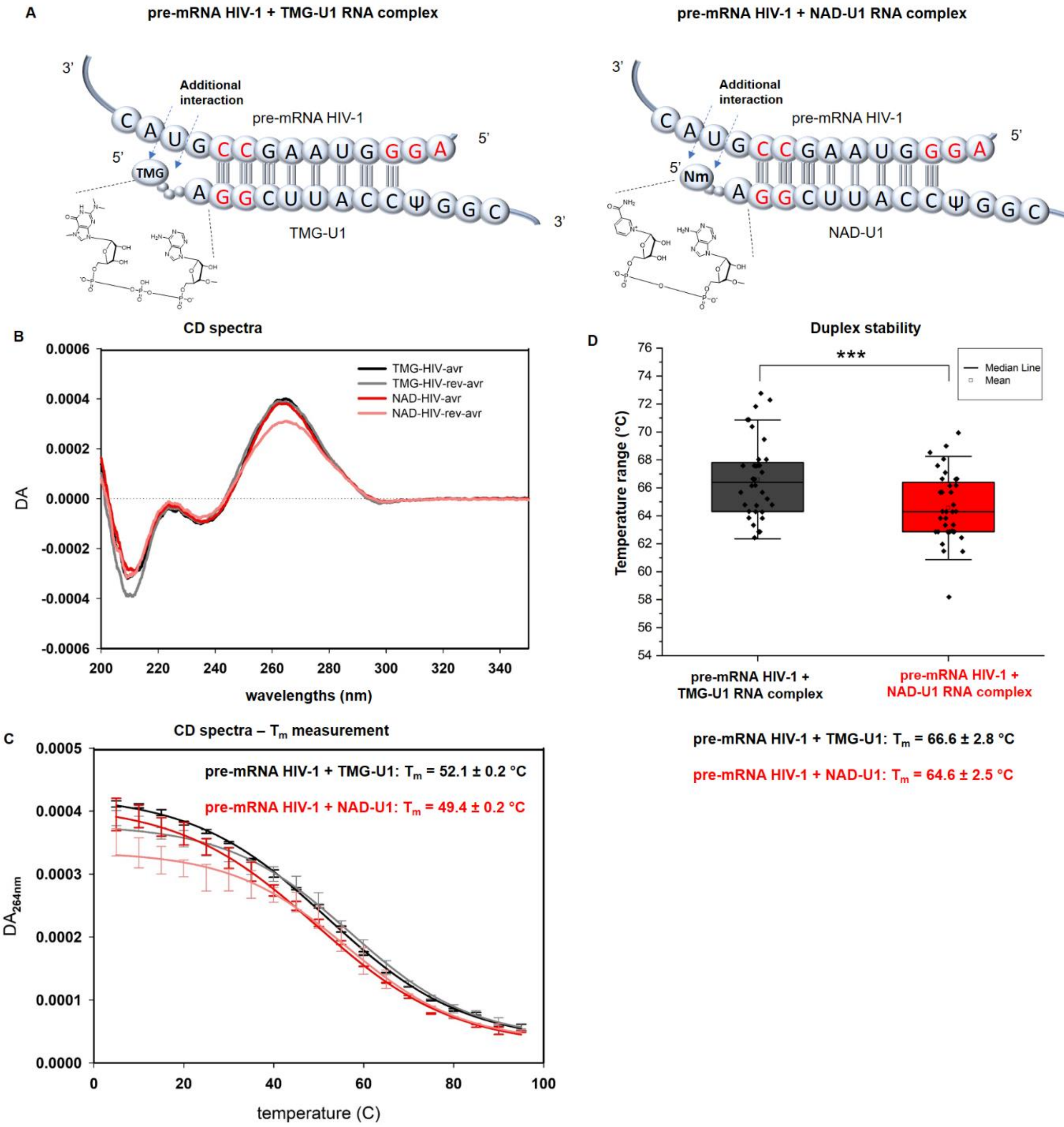


*Figure S 7:* ***NAD cap of U1 snRNA (without pseudouridines) destabilizes the complex with HIV–1 pre–mRNA.*** *(A) Scheme of complex formed by complemental regions of pre–mRNA HIV–1 with either TMG–U1 (left) or NAD–U1 (right). (B) CD spectra of HIV–1 pre–mRNA with TMG–U1 (black or dark grey) and HIV–1 pre–mRNA with NAD–U1 (red and pink). (C) Tm measurement of duplex stability employing CD. (D) Duplex stability measurement of pre–mRNA HIV–1 with TMG–U1 (dark grey) or NAD–U1 (red) by light cycler.*

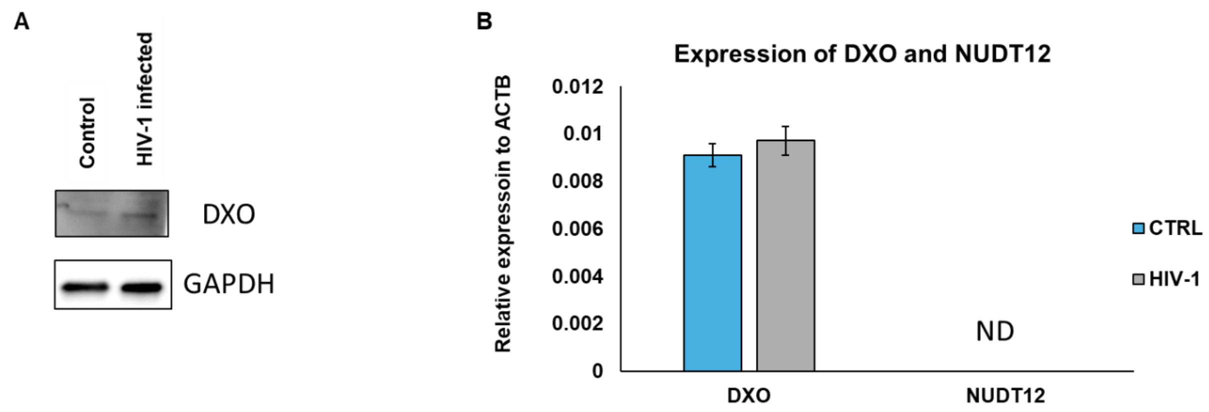


*Figure S 8: Expression of DXO and Nudt12 in control and HIV–1 infected cells. (A) Western blot analysis of DXO and GAPDH (as control) in control cells and HIV–1 infected cells. (B) RT–qPCR of DXO and Nudt12 in control and HIV–1 infected cells.*

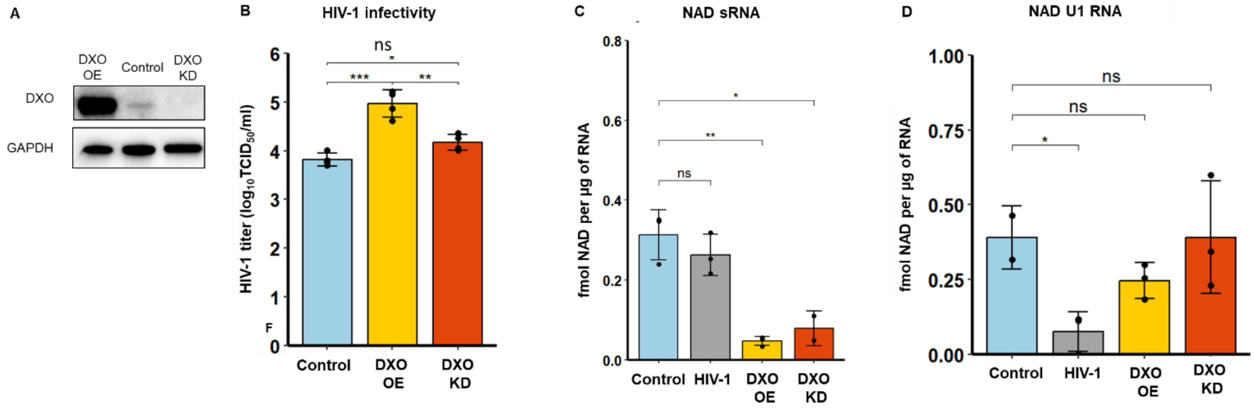


*Figure S 9:* ***Decrease of NAD RNA capping leads to increased HIV–1 infectivity.*** *(A) Western blot analysis of DXO and GAPDH (as a control) in control cells and cells with overexpressed (DXO OE) or downregulated DXO (DXO KD). (B) HIV–1 infectivity determined in control cells and cells with overexpressed (DXO OE) or downregulated (DXO KD) DXO (ANOVA test). (C LC–MS quantification of the NAD RNA cap level in sRNA isolated from control cells and cells with overexpressed (DXO OE) or downregulated (DXO KD) DXO (t–test). (D) LC–MS quantification of the NAD RNA cap level in pulled down U1 RNA (t–test). All the experiments were performed in biological triplicates.*

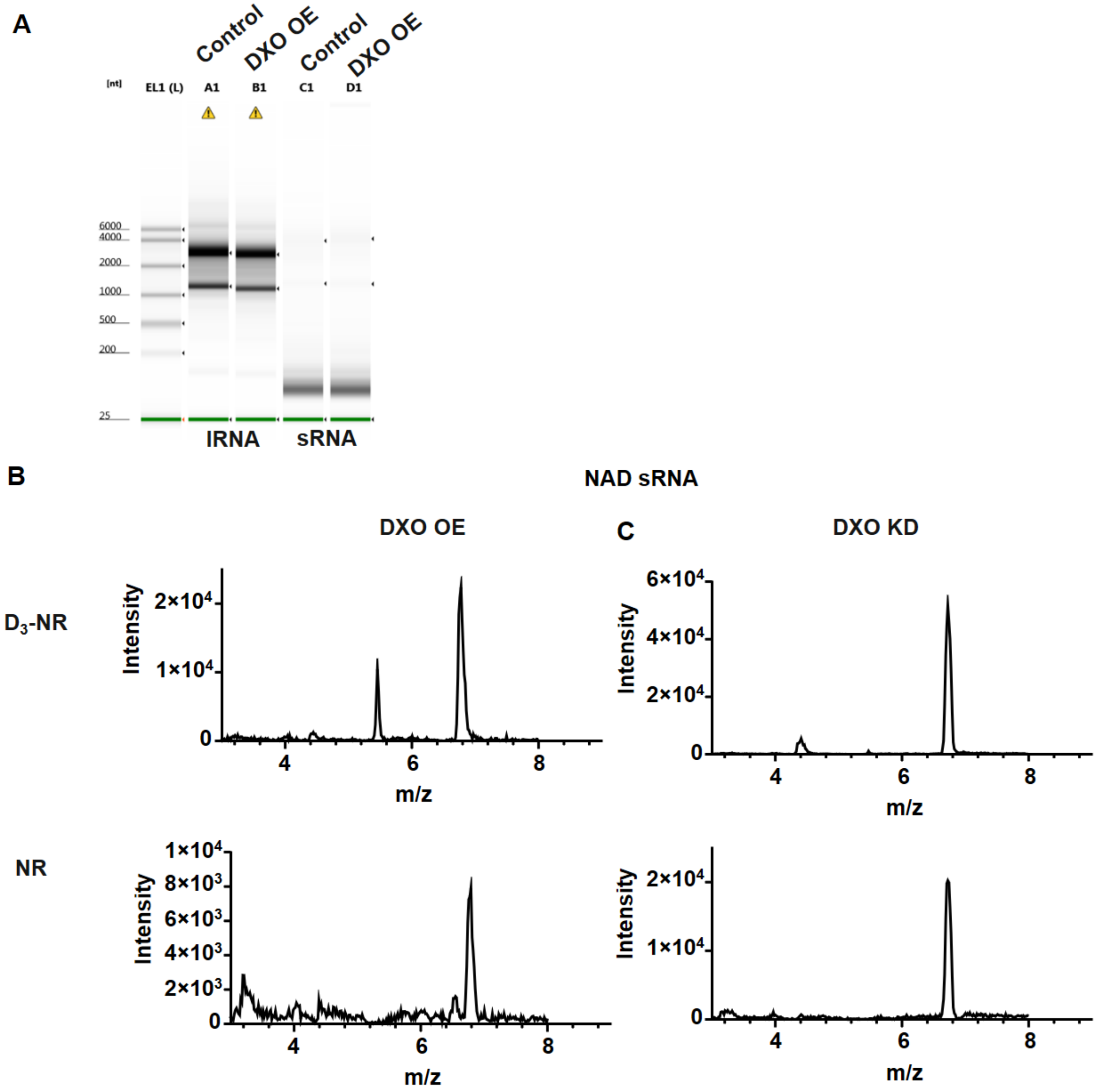


*Figure S 10:* ***LC–MS analysis of isolated sRNA from MT4–DXO OE and MT4–DXO KD cells.*** *(A) Example of sRNA and long RNA (lRNA) franctionation from control and MT4–DXO OE cells. (B) Extracted ion chromatograms of Tof–MRM transitions 258 →126 for $D_3$–NR used as internal standard and 255→123 for analysed NR in RNA from MT4–DXO OE. (C) Extracted ion chromatograms of Tof–MRM transitions 258 →126 for $D_3$–NR used as internal standard and 255→123 for analysed NR in RNA from MT4–DXO KD cells.*

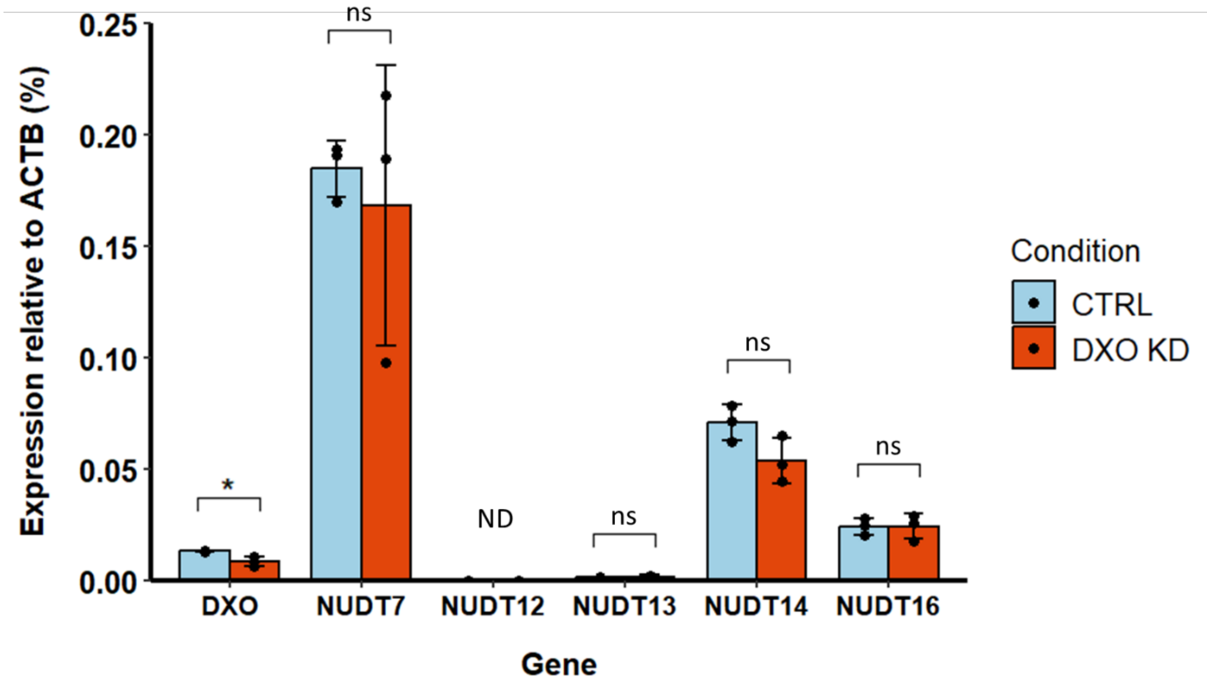


*FIgure S 11:* ***RT–qPCR of DXO and some selected NudiXes in control and DXO KD cells.*** *Experiment was performed in biological triplicate and was normalized to ACTB (actin biotin) mRNA.*

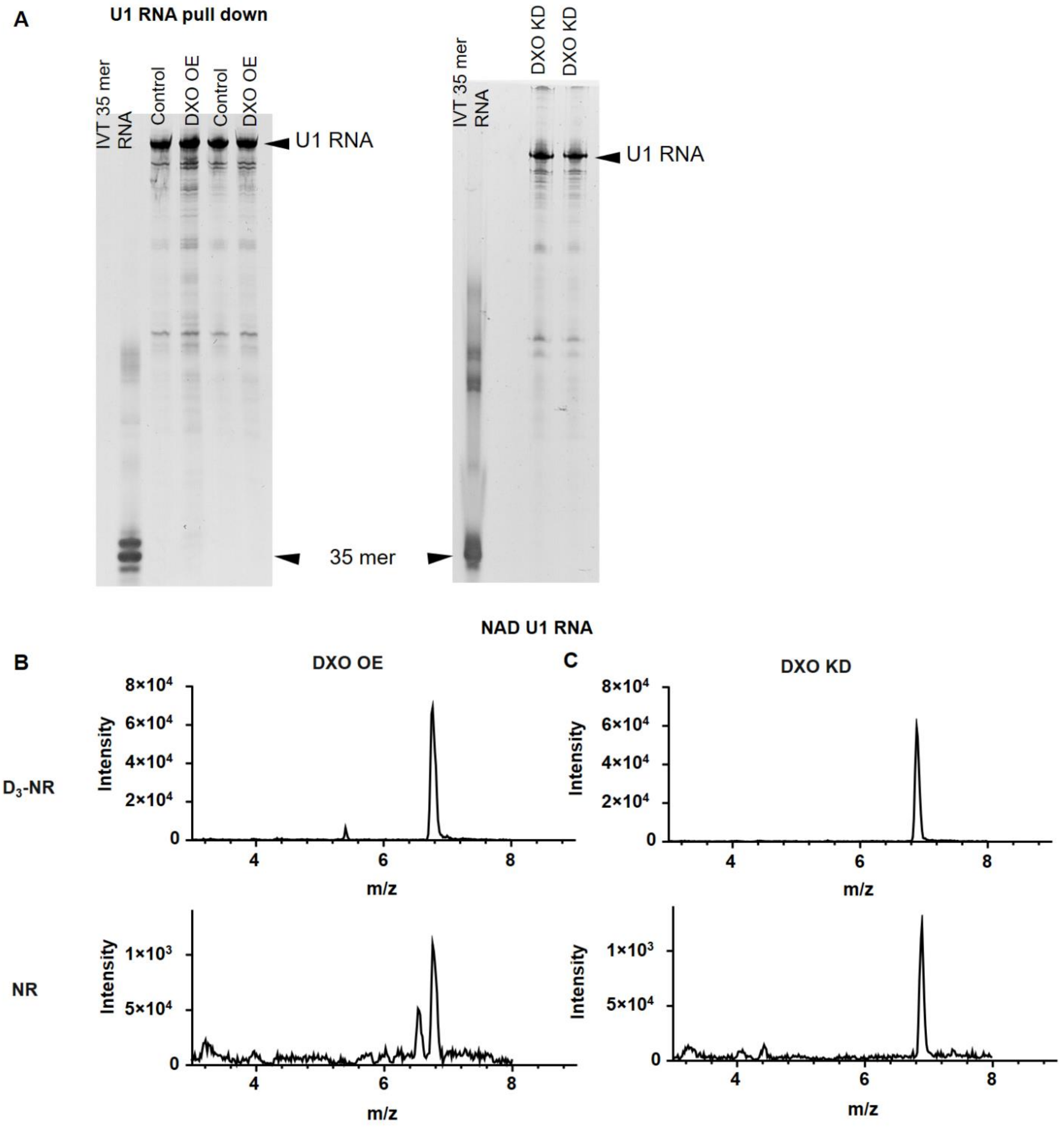


*Figure S 12:* ***LC–MS analysis of isolated U1 RNA pulled down from control and HIV–1 infected cells.*** *(A) PAGE (12.5%) analysis of U1 RNA pulled down from control, MT4–DXO OE and MT4 DXO KD cells. (B) Extracted ion chromatograms of Tof–MRM transitions 258 →126 for $D_3$–NR used as internal standard and 255→123 for analysed NR in U1 RNA from MT4–DXO OE cells. (C) Extracted ion chromatograms of Tof–MRM transitions 258 →126 for $D_3$–NR used as internal standard and 255→123 for analysed NR in U1 RNA from MT4–DXO KD cells.*

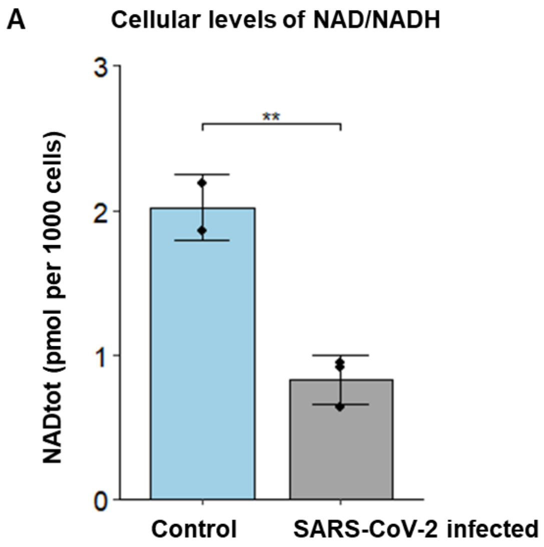


*Figure S 13:* ***Levels of free NAD/NADH*** *(A) in control Vero cells and SARS–CoV–2 infected cells (measured in biological triplicates and technical duplicates, t–test). (B) in control VeroE6 cells and HSV–1 infected cells (measured in biological triplicates and technical duplicates, t–test).*

**Supplementary protocol.**

*Cell culture.*

The human CD4+ T–cell line MT–4 (NIH AIDS Reagent Program, Division of AIDS, NIAID, NIH from Dr. Douglas Richman) and the human embryonic kidney HEK293T cell line (American Type Culture Collection, LGC Standards, UK) were cultured under standard conditions at 37 °C under a humidified (>90%) atmosphere of 5% CO2/95% air. The MT–4 cell line was cultured in RPMI 1640 with stable Glutamine and 25mM Hepes (LM–R1638/500, Biosera) supplemented with 10 % (v/v) FBS (FBS–11A, Capricorn), 1 % (v/v) of Penicillin: Streptomycin solution 1x (L0022, Biowest). The HEK293T cell line was cultured in DMEM with 4.5 g $L^{-1}$ glucose, with stable glutamine, with sodium pyruvate (L0103, Biowest) supplemented with 10% (v/v) FBS 1 % (v/v) of Penicillin: Streptomycin solution 1x. MT–4 cells with and without lentiviral plasmids were harvested, divided to aliquots of 100 $10^6$ cells and collected by centrifugation (225× g, 5 min, at 20 °C). Cells pellets were lysed with 12 ml of RNAzol reagent (R4533, Sigma–Aldrich) and stored at -80°C for further RNA isolation.

*HIV–1 culture.*

MT–4 cells were infected with a cell–free HIV–1 strain NL4–3, which was generated by transient transfection of HEK293T cells with a pNL4–3 plasmid (obtained through NIH AIDS Reagent Program, Division of AIDS, NIAID, NIH from Dr. Malcolm Martin). The infected cultures were subsequently expanded by co–cultivation: 48 h post–infection, cell culture supernatants containing viral particles and infected cells were added to uninfected MT–4 cells ($5\times10^5$ cells per mL) at a ratio of 1:9. The co–culture was synchronized by three successive additions of infected culture supernatant to uninfected MT–4 cells ($5\times 10^5$ cells per mL, the ratio of 1:9, 27 h interval). For the NAD repletion, the medium was supplemented with 10 mM of nicotinamide (NAm, Sigma–Aldrich), the NAD precursor [1, 2].The cell cultures were incubated for additional 40 hours and then harvested.MT–4 cells infected and noninfected were collected by centrifugation (225× g, 5 min, at 20 °C). Cells pellets were washed with PBS and cells were lysed with RNAzol reagent (Sigma–Aldrich). Virus–containing supernatants were filtered (0.45 µm pore size cellulose–acetate filter (VWR)), and stored at −80 °C.HIV–1 titer was measured by 10–fold serial infection of TZM–bl cells (NIH AIDS Reagent Program, Division of AIDS, NIAID, NIH from Dr. John C. Kappes and Dr. Xiaoyun Wu) in triplicate, calculated by Reed–Muench method and expressed as 50% tissue culture infectious dose.

*Viral particles purification via Sucrose cushion isolation*

Virus particles were concentrated from cleared culture medium by centrifugation through a cushion of 20% (w/w) sucrose in phosphate–buffered saline (PBS) (90000 × g, 90 min, 4 °C). The pellet was resuspended in RNase/DNase buffer (Tris–HCl 100 mM, $MgCl_2$ 25 mM, $CaCl_2$ 25 mM) with DNase I (10 U $mL^{-1}$, New England BioLabs – NEB), RNase I (200 U $mL^{-1}$) and RNase A (20 mg $mL^{-1}$, both ThermoFisher Scientific) and incubated 2 h at 37 °C. RNase/DNase treatment was stopped by adding RNAzol (Sigma–Aldrich).

*NAD/NADH measurement.*

NAD total levels in MT–4 cells and MT–4 cells infected by HIV–1 were assessed using the NAD/NADH Quantification Kit (Sigma–Aldrich) according to the manufacturer's protocol. $2\times10^5$ MT–4 cells from each set were pelleted and the extract together with NADH standards (all in duplicates) were measured in a 96 well flat–bottom plate in three replicates. To evaluate the efficiency of NAD repletion, eight differently prepared MT–4 cell lines (MT–4, MT–4 infected by HIV–1, MT–4 with DXO OE, MT–4 with DXO OE and infected by HIV–1, all conditions with or without NAD repletion) were measured as described and expressed as fold–increase of NAD levels.

*Isolation of small RNA fractions from the cell culture.*

Short RNA fraction was purified according to the RNAzol manufacturer's protocol followed by washing with urea solution to remove non–covalently bound NAD [3]. First, sRNA was washed thrice with 8.3 M urea, then twice with RNAse–free water, twice with 4.15 M urea and 4 times with RNAse–free water. The RNA concentration was determined on NanoDrop ONE (ThermoFisher Scientific) and the RNA sample quality control was performed on a 4200 TapeStation System (Agilent).

*NAD captureSeq library preparation.*

NAD captureSeq libraries from four biological replicates for each sample were performed as described[4]. Briefly, only short fraction of RNA from MT–4 cells infected by HIV–1 and MT–4 cells noninfected were used for the library preparation. The most critical step is the enzyme reaction catalyzed by ADP–ribosylcyclase (ADPRC, Sigma–Aldrich), which is an enzyme specific only for NAD–capped RNA but inactive on canonical RNA. Samples not treated by ADPRC were used as a negative control for non–specifically bound RNA. 100 µg of sRNA for each sample was incubated with 4–pentyn–1–ol (10% (v/v), Sigma–Aldrich) and ADPRC (2.5 µg) in 50 mM Na–HEPES (pH 7.0, Sigma–Aldrich) and 5 mM $MgCl_2$ (Sigma–Aldrich) for 30 min at 37 °C. The reaction was stopped by phenol/ether (Roti–Aqua–Phenol: Carl Roth, Diethyl ether: Penta) extraction and RNA was ethanol precipitated. NAD–capped RNA was then biotinylated (biotin–PEG3–azide, 250 mM, Jena Bioscience) via a copper–catalyzed azide–alkyne cycloaddition (CuAAC) in a freshly prepared mixture of CuSO4 (1 mM, Sigma–Aldrich), THPTA (0.5 mM, Sigma–Aldrich) and sodium ascorbate (2 mM, Sigma–Aldrich) for 30 min at 25 °C, with shaking at 350 r.p.m. The biotinylated RNA was purified by phenol/ether extraction and ethanol precipitated. Mobicol Classic columns (MoBiTec, GmbH) were assembled and 50 µL of streptavidin Sepharose (GE Healthcare) was transferred to each column. After washing the beads by adding three times 200 µL of immobilization buffer (10 mM Na–HEPES, 1 M NaCl and 5 mM EDTA, pH 7.2) to each column and centrifuge at ≥16,100 g for 1 min at rt, the beads were blocked with acetylated BSA (100 µg $mL^{-1}$, Sigma–Aldrich) in 100 µL of immobilization buffer for 20 min at 20 °C, with shaking at 1000 r.p.m. After three more washes, the RNA was immobilized on the beads for 1 h at 20 °C, with shaking at 1000 r.p.m. The beads were washed five times with 200 µL of streptavidin wash buffer (50 mM Tris–HCl and 8 M urea, pH 7.4, both Sigma–Aldrich), then equilibrated by washing three times with 200 µL of 1x standard ligation buffer (50 mM Tris–HCl, 10 mM $MgCl_2$, pH 7.4), blocked with acetylated BSA, and washed three more times with 1x standard ligation buffer as described previously. 30 µL of ligation mixture containing adenylated RNA 3'adaptor (5 µM), 1x standard ligation buffer, DMSO (15%, Sigma–Aldrich), acetylated BSA (1.5 µg), 2–mercaptoethanol (50 mM, Sigma–Aldrich), T4 RNA ligase (15 U, Thermo Fisher Scientific), and T4 RNA ligase 2, truncated K227Q (300 U, New England BioLabs) was added to the beads and the mixture was incubate at 4 °C overnight (≥16 h). The biotinylated RNA was rebound by adding 7.5 µL 5 M NaCl and incubated for 1 hour at 20 °C, with shaking at 1,000 r.p.m. After washing the beads five times with the streptavidin wash buffer, they were equilibrated by washing three times with 1× first–strand buffer (50 mM Tris–HCl, 75 mM KCl and 5 mM $MgCl_2$, pH 8.3), blocked with acetylated BSA, and washed another three times with 1× first–strand buffer. The beads were covered with 30 µL of reverse transcription mixture containing dNTPs (0.5 mM each, New England BioLabs), acetylated BSA (1,5 µg), DTT (5 mM), 1x first strand buffer, reverse primer (5 µM), and Superscript III reverse transcriptase (300 U, Thermo Fisher Scientific) and incubated for 1 hour at 37 °C. The rebinding procedure was repeated and samples were washed five times with streptavidin wash buffer and three times with 1x ExoI buffer (Thermo Fisher Scientific). The beads were blocked with acetylated BSA in ExoI buffer and again washed three times with 1x ExoI buffer. The free primers were digested by adding 30 µL of 1x ExoI buffer and 30 U of ExoI (Thermo Fisher Scientific) and the mixture was incubated for 30 min at 37 °C. The beads were washed five times with streptavidin wash buffer and three times with immobilization buffer. The RNA was digested by adding 100 µl of 0.15 M NaOH solution and incubating for 25 min at 55 °C. The columns were centrifuged and washed with 100 µl of $H_2O$. The flow–through was ethanol precipitated and the cDNA pellet was dissolved in 19 µl of Terminal deoxynucleotidyl transferase (TdT) tailing mixture containing 1x TdT buffer (Thermo Fisher Scientific) and CTP (1.25 mM). After adding 20 U of TdT (Thermo Fisher Scientific), the mixture was incubated in thermocycler for 30 min at 37 °C and after the enzyme was thermally denatured by heating to 70 °C for 10 min. Then, the second adaptor was ligated on the cDNA by adding 60 µl of reaction mixture containing 1x standard ligation buffer, cDNA anchor (5 µM each strand), ATP (10 µM), and T4 DNA ligase (120 Weiss U, Thermo Fisher Scientific) and incubating for 16 h at 4 °C. PCR amplification was performed with barcoded reverse PCR primers in 25 cycles in Taq DNA Pol reaction buffer 1 × (Thermo Fisher Scientific), with 1 µM of each reverse barcoded primer and forward primer (Table SI1), 0.2 mM dNTPs (each) and 2.5 U of DreamTaq DNA Polymerase (Thermo Fisher Scientific) in 50 µL of total reaction mixture. Initial denaturation was performed at 95 °C for 3 min, following by annealing for 30 s at 68 °C, elongation for 30 s at 72 °C and denaturation for 30 s at 95 °C. Final extension was performed at 72 °C for 5 min. PCR reaction mixture was loaded on 2.5% agarose gel (140 V for 2 h) and post–stained with SYBR gold (Thermo Fisher Scientific). Fractions between 100–400 nt were cut and DNA was extracted from the gel by Monarch DNA Gel extraction Kit (NEB). Multiplexed samples were submitted to the IMG Genomics and Bioinformatics facility for library synthesis using the NextSeq® 500/550 High Output Kit v2, 75 cycles (Illumina) and sequencing on a NextSeq 500/550, Illumina.

*Small RNA–seq library preparation.*

To verify the amounts of sRNAs in samples, three biological replicates of sRNA from MT–4 cells and MT–4 cells infected by HIV–1 were submitted for miRNA (LP–170) library preparation to SEQme. The library synthesis was done using the NEBNext®Small RNA Library Prep Set for Illumina with NEBNext Multiplex Oligos for Illumina (Index Primers Set 1–3) and sequenced on a NovaSeq 6000 System (SE 50 bp, 350–400 millions of reads, DS–270).

*Data processing and bioinformatic analysis.*

Adapters and low–quality bases were trimmed using TrimGalore! v.0.4.1 (https://github.com/FelixKrueger/TrimGalore). Because of a bias at the ends of the reads, additional 5 bases were trimmed from both 5' and 3' end of the reads. Reads were mapped to the human GRCh38 genome and HIV–1 HIVNL43 NC_001802.1 genome using Hisat2 v.2.0.1. To analyse NAD presence at HIV RNA cap, we quantified RPM in the first 200 bp of HIV transcribed region and quantified the enrichment of ADPC–treated samples over controls using Seqmonk v1.47.2 (https://github.com/s-andrews/SeqMonk). Enrichment of human shortRNAs was quantified as RPM in ADPRC–treated over control samples using Seqmonk v1.47.2. To select the best candidate shortRNAs with depletion or increase of NAD–cap abundance after HIV infection, we applied a set of criteria removing shortRNAs with low read count and/or low enrichment over control (-ADPRC): (I) raw read count has to be 2 or higher for at least 2 replicates of at least one sample type (+/-ADPRC +/- HIV infection), (II) enrichment over control higher than 2 in at least one replicate and enrichment above 1 in at least one additional replicate in either HIV infected or non–infected samples, (III) of all 4 replicates, 2 or more have to show depletion/increase in the same direction (difference between HIV non–infected and infected above 0.2), and neither in the opposite direction, and maximum 2 replicates the lack of enrichment (enrichment <1), (IV) total raw read count in combined replicates of +ADPRC samples (either non–infected or infected) above 20, (V) the number of reads in +ADPRC samples above 5 in the replicates showing enrichment above 1. Differential expression of sRNAs of all sRNA classes annotated within Seqmonk v1.47.2 (GRCh38v100 genome annotation) was analysed using DESeq2 implemented within Seqmonk v1.47.2. Splicing efficiency of HIV–1 transcript was analysed as ratio of read count in random regions within introns to read count in size matched random regions in exon 3. Sequences of main donor and acceptor splice sites in HIV–1 main splicing variants (according to [5]) were obtained from [6] and matched against HIV–1 HIVNL43 NC_001802.1 genome to generate genomic coordinates of exons and introns. Splicing efficiency of cellular transcripts was quantified as ratio of reads in introns to reads in exons for 1000 genes with the highest expression in combined replicates of untreated MT–4 cells quantified using RNA–seq quantitation pipeline within Seqmonk v1.47.2.

*Deep sequencing data validation through RT–qPCR.*

Real–time PCR was performed to verify the NAD captureSeq data. Three biological replicates of sRNA from infected and noninfected MT–4 cells and cDNA samples after NAD captureSeq were measured in two technical repeats on LightCycler 480 II (Roche) by Luna Universal One–Step RT–qPCR Kit (New England BioLabs) according to the manufacturer protocol. Briefly, 20 µl reaction mix consisting of buffer, enzyme mix, forward and reverse primer (0.4 µM, Table SI1) and 50–100 ng template was prepared. PCR was conducted with the following program: reverse transcription 55 °C for 10 min, initial denaturation at 95 °C for 60 s, followed by 40 cycles of 95 °C for 10 s and 53 °C for 30 s. Finally, a melting curve was performed from 37 °C to 95 °C. The Cp values were calculated by LightCycler 480 Software and the reciprocal values were plotted.

*Preparation of RNA samples for LC–MS analysis*

Short RNA fraction (≈50 µg) or U1 RNA (≈5 µg) was mixed with 20 pmol of NudC pyrophosphatase (New England Biolabs, #M0607S) and 1 U of shrimp alkaline phosphatase (New England Biolabs, #M0371S) in buffer containing 50 µM ammonium formate (pH 7.0, Sigma–aldrich, #09735), 10 mM DTT (shipped with NudC) and 10 mM $MgCl_2$ (Sigma–aldrich, #63068) and molecular biology grade water (Sigma–aldrich, #W4502). Total reaction volume was 40 µL. Mixture was incubated for 30 min at 37 °C. Right after, 10 µL of 1.5 nM nicotine amide riboside–d4 internal standard (Torronto Research Chemicals, #TRC–N407772) was spiked to the sample and the mixture was filtered through 10 kDa Vivacon 500 filter (Sartorius, #VN01H02) at 14 000× g for 15 min at 10°C. Filtrate was immediately mixed with 2 equivalents of acetonitrile (LC–MS grade) in HPLC vial and analysed by LC–MS.

*LC–MS analysis*

Samples were separated with HPLC (Acquity H–class, Waters) equipped with Xbridge Premier BEH amide column (2.5 µm, 4.6 mm X 150 mm, Waters) heated to 35°C. Autosampler was kept at 10°C , injection volume was 50 µL and the flow rate was 1 mL/min. Mobile phase A contained 10 mM ammonium formate (pH 9.0, Fisher, #A11550) in 90% (v/v) acetonitrile (Fisher, #A955212) and mobile phase B 10 mM ammonium formate (pH 9.0) in ultrapure water (18.2 MΩ.cm, Purelab Chorus system, Elga). The gradient of separation was following: 0 min 10% (v/v) B, 2 min 10% B, 8 min 33% B, 10 min 33% B, 10.1 min 10% B, 15 min 10% B. The detection of analytes was performed using Xevo G2–XS QTof mass spectrometer (Waters) equipped with an electrospray ionization source. The spray voltage was 3 kV, sampling cone 20 and source offset 40 V. Source temperature was kept at 150°C and desolvation gas at 500°C. Gas flow rates were 50 L/h for cone gas and 1000 L $h^{-1}$ for desolvation gas. Mass spectrometer was oparated in positive ion mode in 50 – 600 m/z range in sensitive mode with Tof–MRM function and 255→123 transition. Scan time was 1 s and collision gas energy was 3. Nicotine amide riboside was identified based on identical retention time with spiked internal standard and quantified with external calibration curve (spiked sample blanks, which underwent same procedure as analysed RNA) consisting of 4 calibration points (4.6, 14, 42 and 125 pM concentration of nicotine amide riboside) fitted with linear regression. Each point was normalized to isotopically labelled internal standard and is an average of two separate injections.

*RNA in vitro transcription.*
In vitro transcription was performed as described [7] in a 50 µL mixture (0.2 µM of template DNA (Table SI1), 1 mM of each NTP (New England BioLabs), 5% dimethyl sulfoxide (DMSO, New England BioLabs), 0.12% triton X–100 (Sigma–Aldrich), 10 mM dithiothreitol (DTT), 4.8 mM $MgCl_2$ (Sigma–Aldrich), 1× reaction buffer for T7 RNAP (New England BioLabs) and 125 U of T7 RNAP (New England BioLabs). For capped RNA, 8 mM of NAD (Sigma–Aldrich) or TMG cap was added. For the production of 20 nt long U1 RNA containing pseudouridine, the UTP was replaced with pseudo–UTP (Jena Bioscience). The mixture was incubated for 2 h at 37 °C. For higher yield, T7 RNAP was added again into the mixture and incubated for additional two hours. The DNA template was digested by DNAse I (New England BioLabs) at 37 °C for 45 min and the enzyme was heat inactivated at 75 °C for 10 min. To obtain only capped RNA, the samples were treated with 5′–polyphosphatase (Epicenter) for 30 min at 37 °C and then with Terminator™ 5′–phosphate–dependent exonuclease (Epicenter) for 1 h at 30 °C. Between each step, samples were purified using Clean and concentrator (Zymo).

*Assessment of melting temperature by circular dichroism.*
The ECD spectra were measured on a Jasco 815 spectropolarimeter (Tokyo, Japan) equipped with the Peltier type temperature control system PTC–423S/L at room temperature in spectral range from 200 nm to 350 nm, using a 0.2 cm path length with standard instrument sensitivity, with the scanning speed of 10nm $min^{-1}$, response time of 8 s, and 2 spectra accumulations. For measurements 26000 ng of RNA duplex with TMG–U1 RNA and/or RNA duplex with NAD–U1 was dissolved in annealing buffer (10 mM Tris, 50 mM NaCl, 1 mM EDTA pH 7.8). The final spectra were expressed as differential absorption $\Delta$A. Melting temperature was obtained as results of measurement of CD signal at 264 nm in temperature range from 5 °C to 95 °C with temperature slope 24 °C/h, step 1 °C and time constatnt 8 sec. Each measurement was repeated 3 times. Melting temperature was calculated using sigmoid fitting by program Sigmaplot 12.5 (Systat software).

*U1 RNA and HIV–1 mRNA complex stability assay.*
20 nt long U1 RNA (capped either with TMG or NAD cap) and HIV–1 mRNA (Table SI1) were prepared by in vitro transcription. 150 ng of U1 RNA (TMG or NAD cap) and 150 ng of HIV–1 mRNA were mixed with ResoLight dye (1x, Roche) and annealing buffer (10 mM Tris, 50 mM NaCl, 1 mM EDTA pH 7.8). Three technical replicates were prepared three times and measured on LightCycler 480 II (Roche). High–resolution melting curves were obtained by measuring the complex stability by temperature increase to 80 °C and then decrease to 20 °C with a ramp rate 0.01 °C/s in three cycles and thus obtaining a total of 27 melting curves per sample. HRM curve analysis was performed using the LightCycler 480 Software.

*Radioactively labelled U1 RNA preparation.*
DNA template for U1 RNA (164 nt full length) was prepared from plasmid (U1–SP65 plasmid containing the main U1 variant RNA–U1–1) via PCR in 50 µL reaction mix consisting of DreamTaq buffer (1x, Thermo

Fisher Scientific), DreamTaq DNA polymerase (1.25 U, Thermo Fisher Scientific), forward and reverse primer (0.5 µM, Table SI1), dNTPS (2mM each, New England BioLabs) and 400 ng template. PCR was conducted with the following program: initial denaturation at 95 °C for 2 min, followed by 30 cycles of 95 °C for 30 s, 60 °C for 30 s and 72 °C 1 min, the final extension was performed at 72 °C for 5 min. PCR reaction mixture was loaded on 1% agarose gel (140 V for 2 h) and post–stained with SYBR gold (Thermo Fisher Scientific). The product bend was cut and DNA was extracted from the gel by Monarch DNA Gel extraction Kit (NEB). $^{32}$P–GTP labelled U1 RNA was prepared by in vitro transcription as described previously with addition of 0.5 µL of α–$^{32}$P–GTP (3.3 µM, 10 µCi µL$^{-1}$; Hartmann analytic), with either TMG or NAD cap.

*U1 RNA decapping by DXO*

1000 ng of $^{32}$P–GTP labelled U1 RNA was incubated with the DXO decapping enzyme at a final concentration of 50 nM in a total volume of 25 µl in decapping buffer (10 mM Tris pH 7.5, 100 mM KCl, 2 mM DTT, 2 mM $MgCl_2$ and 2 mM $MnCl_2$) at 37 °C. Reactions were stopped by the addition of RNA loading dye (Thermo Fisher Scientific) after 0, 10, 30, 60, 120, and 240 minutes. 20 µl from each sample was then loaded in per well. The denaturing polyacrylamide gel (8%) was prepared from Rotiphorese gel (Carl Roth) in 1x TBE buffer and supplemented with 7 M urea, acryloylaminophenylboronic acid (APB, 0.4% (w/v)) 48 and 20 µl of Tetramethylethylenediamine (TEMED). The polymerized gel was pre–run at 600 V for 30 min in 1x TBE buffer and then run at 600 V for 4.5 hours. The gel was incubated with phosphor imaging plate for 45 minutes (GE healthcare), scanned using Typhoon FLA 9500 (GE Healthcare), and analysed using ImageJ.

*U1 RNA pull down.*

Annealing of 1 nmol of biotinylated U1 probe (Table SI1) to 150–300 µg of sRNA was performed in mixture supplemented with 10 mM Tris, 0.9 M TMAC and 100 mM EDTA, pH 7.8, by a temperature slowly decreasing from 65 °C to 25 °C. Streptavidin Sepharose™ High Performance (Cytiva) were prepared based on manufacturer protocol. Briefly, the beads were resuspended and 20 µL was transferred to a Mobicol column and washed twice with 1x PBS and once with 1x RNA pull–down buffer (10 mM Tris–HCl, 100 mM EDTA, 0.9 M TMAC, pH 7.6). To keep solutions RNAse–free, all buffers were DEPC–treated. After washing, the beads were resuspended in annealing mixture. Samples were incubated for 10 mins at 25 °C using gentle rotation. The beads were washed 6 times with 10 mM Tris (pH 7.6) and resuspended in RNase–free water. To dissociate the captured U1 RNA, samples were incubated for 10 minutes at 75°C. This step was repeated twice. The concentration of released U1 RNA was measured on NanoDrop ONE (ThermoFisher Scientific) and the purity of samples was validated via 12.5 % PAGE.

*Preparation of lentiviral vectors for DXO OE and DXO KD.*

HEK293T cells were transfected at 20% confluency in a 100 mm plate with a mixture of two helper plasmids pCMV–VSV–G (4.5 µg) and pCMV–dR82dvpr (4.5 µg) (obtained from the Keckesova lab, IOCB Prague), and 6 µg shDXO or DXO–OE plasmids (see below), using the Lipofectamine 3000 Reagent (ThermoFisher #L3000001). For DXO knock–down, shRNA plasmid from VectorBuilder VB900039–9953mhp (https://en.vectorbuilder.com/vector/VB900039–9953mhp.html, target sequence TAGCTGAGCCTCGGAACAAAC) was used, and the humen DXO gene expression plasmid VB900001–2728fsj, (VectorBuilder, https://en.vectorbuilder.com/vector/VB900001–2728fsj.html) was used to generate DXO OE cell lines. Three days after transfection, 12 mL of cell–free supernatant medium containing the lentiviral particles was harvested, filtered through 0.45 µm filter, and stored at -80 °C until further use.

*Preparation of stable MT4–DXO OE and MT4–DXO KD cell lines.*
For the transduction, 5 million MT4 cells were harvested (225× g, 5 min, at 20 °C) and resuspended in 1 mL of lentivirus containing supernatant. The transduction suspension was incubated for 2 hours at 37 °C, 5% $CO_2$ and afterwards transferred into p25 flask with 5 mL of fresh RPMI media. After 24 hours, the cells were centrifuged (225× g, 5 min, at 20 °C), 5 ml RPMI with 2.25 μg $mL^{-1}$ puromycin (ThermoFisher A11138–03) was added to the flask, resulting in final 1.125 μg/mL puromycin as selection marker. The cells were selected for 7 days with 1.125 μg $mL^{-1}$ puromycine and the medium was subsequently replaced by fresh RPMI (no puromycine). During cell expansion, the maintaining 1 μg $mL^{-1}$ puromycin selection was performed every two weeks for 3 days. The stability of the integration and population purity (>90% manipulated cells) was monitored by FACS using EGFP–T2A–Puro element in the lentiviral vectors and DXO levels were monitored by western blotting.

*Western blot.*
Aliquots of 500000 MT4 control cells, MT4–DXO OE and MT4–DXO KD cells were collected by centrifugation (225× g, 5 min, at 20 °C). Proteins were extracted using RIPA Lysis and Extraction Buffer (Thermo Fisher Scientific #89900), cOmplete™, EDTA–free Protease Inhibitor Cocktail, Sigma Aldrich #04693116001) and the protein concentration in the lysate was measured using DC™ Protein Assay Kit II (Bio–Rad #5000112) on Tecan (Schoeller). 50 µg and 10 µg of protein lysate for DXO respectively GAPDH detection was loaded on the 10–well protein gels (Mini–PROTEAN TGX, Bio–Rad) and run at 120 V. Gels were wet transferred on PVDF membranes (Thermo Fisher Scientific) at 30 V for 2 hours. Primary antibody DOM3Z Rabbit PolyAb (Proteintech Europe #11015–2–AP) diluted 1:500 or GAPDH (14C10) Rabbit mAb (Cell Signaling Technology #2118) diluted 1:5000 was incubated with the membrane overnight at 4 °C. Next, six washes were performed with PBS 0.1% Tween–20 (PBST) for 10 minutes at room temperature before the addition of secondary antibody (1:10.000, peroxidase conjugated AffiniPure Goat Anti rabbit IgG (Jackson Immuno research #111–035–003) for 1 hour at room temperature. Afterwards, six 10 minutes washes were performed with PBS 0.1% Tween–20 (PBST) at room temperature. Blots were developed using SuperSignal™ West Femto Maximum Sensitivity Substrate (ThermoFisher Scientific #34096) and imaged on the Azure Biosystems (model c600).

*RT–qPCR for DXO and Nudix enzymes mRNA*
500 ng of the lRNA fraction of three biological replicates was reverse–transcribed using LunaScript® RT SuperMix Kit (NEB #E3010) according to manufacturer's instructions. The cDNA was diluted 5 times with deionized water. 2 µL of diluted cDNA per well was used per well (performed in technical duplicate) on LightCycler 480 II (Roche). The Luna® Universal qPCR Master Mix (NEB #M3003) was used according to manufacturer's instructions with forward and reverse primer (0.4 µM) listed in Table SI1. PCR was conducted with the following program: initial denaturation at 95 °C for 60 s, followed by 45 cycles of 95 °C for 10 s and 60 °C for 30 s. Finally, a melting curve analysis was performed from 37 °C to 95 °C. The melting curve analysis and Cq values were calculated by LightCycler 480 Software.

*HIV–1 infectivity determination in DXO–transduced cells.*
The HIV–1 infectivity and NAD repletion in DXO–transduced cells was determined by measuring HIV–1–induced cytopathic effect by XTT cell proliferation assay. Briefly, wild type MT–4 and DXO–transduced MT–4 cells were seeded at 3 $10^4$ cells per well in a 96–well plate in RPMI medium (phenol red–free RPMI, supplemented with 10% FBS, penicillin (100 U $mL^{-1}$), streptomycin (100 µg $mL^{-1}$), 2 mM L–glutamine and 4 mM HEPES, all Sigma Aldrich) with and without 10 mM NAm, infected in triplicate by two–fold serially diluted HIV–1 and cultured at 37 °C and 5% $CO_2$. After 4 days, 50:1 mixture of XTT salt and PMS electron–

coupling reagent (both VWR) was added to the wells and incubated for 4 h at 37 °C in 5% $CO_2$. Formation of orange formazan dye was measured in EnVision plate reader (Perkin Elmer) at 560 nm. HIV–1 titer was calculated by Reed–Muench method and expressed as 50% tissue culture infectious dose.

*Cell cultures used for NAD/NADH measurement*

Vero cells and VeroE6 cells (American Type Culture Collection) were cultured under standard conditions at 37 °C under a humidified (>90%) atmosphere of 5% CO2/95% air. The lines were cultured in DMEM with 4.5 g $L^{-1}$ glucose, with stable glutamine, with sodium pyruvate (L0103, Biowest) supplemented with 10% (v/v) FBS 1% (v/v) of Penicillin: Streptomycin solution 1x. $2\times10^5$ cells were then seeded in 24 well plates and infected by 100 µl of the diluted virus (HSV–1 BSA stabil. MII IIE21 or HCoV–229E virus VR–740 from ATCC) at TCID50 $10^5$,99 IU $mL^{-1}$. The NAD/NADH content was measured after 24 hours post infection based on the same protocol as the MT4 cells.

*Synthesis of $m_3^{2,2,7}$GpppApG (TMGpppApG, TMG cap).*

To enable synthesis of highly homogenous 5'–TMG–capped RNA by in vitro transcription we designed a trinucleotide TMG cap analog (TMGpppApG or $m_3^{2,2,7}$GpppApG) following the synthetic pathway described previously for NAD–derived trinucleotides.[8] All organic solvents that were used in the chemical syntheses under anhydrous conditions: dimethyl sulfoxide (DMSO, Honeywell, HPLC grade), N, N–dimethylformamide (DMF, anhydrous, Sigma–Aldrich) and acetonitrile (ACN, HPLC grade, J.T.Baker), were additionally dried over 4A molecular sieves. Other solvents: diethyl ether (CHEMPUR, p.a.), acetone (CHEMPUR, p.a.), methanol (MeOH, J.T. Baker, HPLC grade), acetic anhydride (CHEMPUR, p.a.) were used as received. Reversed phase HPLC (RP–HPLC) was carried out on a Agilent system for the analysis and final purification of the compounds. A Gemini column (NX–C18, 150 mm x 4.6 mm, 3 µm) was used to analyse the reactions progress with a flow rate of 1 mL $min^{-1}$. The solution of 50 mM ammonium acetate ($CH_3COONH_4$) pH 5.9, and the mixture of 50 mM $CH_3COONH_4$, pH 5.9 and ACN (1/1, V/V) were used as buffers. HiCHROM C18, 150 mm x10 mm, 5 µm, with a flow rate of 4.7 mL $min^{-1}$ was used as semi–preparative column. Buffers for RP–HPLC were as follows: A: 50 mM $CH_3COONH_4$, pH 5.9, B: 50 mM $CH_3COONH_4$, pH 5.9 / MeOH, 1/1, V/V. DEAE Sephadex A–25 was used for purification by ion exchange column chromatography. Triethylammonium bicarbonate (TEAB) buffer in deionized water was used as the mobile phase in a linear gradient: 0 to 0.7 M TEAB for nucleoside monophosphate, 0 to 0.9 M TEAB for nucleoside diphosphate, 0 to 1.2 M TEAB for dinucleoside triphosphate.

TMG cap ($m_3^{2,2,7}$GpppApG) was synthesised following the procedures formerly reported with our modifications[8, 9]. The synthetic pathway consisted of 9 steps. A convergent synthesis approach was used to minimize the loss of yields in each step. The starting substrate was commercially available guanosine. The scale of the individual steps as well as the techniques used to purify the intermediates were tailored to our needs. Guanosine methylation at the N7 position was carried out in the final step of the synthesis of $m_3^{2,2,7}$GDP to avoid degradation of the compound during the 5'–phosphorylation.

a. 2',3',5'–tri–*O*–acetylguanosine **(1)**

Guanosine (8.83 mmol, 2.5 g, Carbosynth) was suspended in anhydrous acetonitrile (100 mL) and then 4–dimethylaminopyridine (DMAP, 0.663 mmol, 81 mg, Sigma–Aldrich) and TEA (4.87 mL) were added. The solution was stirred for 5 minutes on a magnetic stirrer at room temperature and then acetic anhydride (3 mL) was added. The reaction was carried out with continuous stirring for 50 minutes at room temperature in a closed flask. After this time, the reaction was terminated by adding 1.25 mL of methanol to the flask and stirring for a while more. The solvents were evaporated to dryness on a rotary evaporator and then cold acetone (50 mL) was added and refrigerated for 0.5h. A white precipitate precipitated on the flask, which was filtered on a Buchner funnel and washed with an additional portion of cold acetone (50 mL). The product was dried in a desiccator to give 2.18 g of the compound as a white powder. The reaction progress was controlled by RP–HPLC and low resolution MS.

b. 2’,3’,5’–Tri–O–acetyl–2–N,2–N–dimethylguanosine **(2)**

To a solution of compound **(1)** (5.0 g, 12 mmol, 1 eq.) in 99.9% acetic acid ($CH_3COOH$, 150 mL, J.T.Baker) paraformaldehyde (1.1 g, 37 mmol, 3 eq., Sigma–Aldrich ) was added and stirred for 3h at 45°C. After this time, sodium cyanoborohydride ($NaBH_3CN$, 2.3 g, 37 mmol, 3 eq., Sigma–Aldrich) was added to the mixture and again stirred vigorously for 3 h at 45°C. The sequence of adding paraformaldehyde and $NaBH_3CN$ at 3 h intervals was repeated two more times, adding a total of 9 eq. of each. The progress of the reaction was monitored by RP–HPLC. The solvent was evaporated on a rotary evaporator and the precipitate was dissolved in a mixture of dichloromethane (DCM / pyridine, 1/1, v/v). The solution was washed 3 times with saturated $NaHCO_3$ and the aqueous layers were extracted again with DCM / pyridine solution (2/1, v/v). The organic layer was collected and dried over $Na_2SO_4$ and then filtered and evaporated on a rotary evaporator with toluene until residual pyridine was removed. The purity of the compound was confirmed by low resolution MS. The product was obtained as a pale grey solid.

c. 2–N,2–N–dimethylguanosine **(3)**

Compound **(2)** (5 g, 12.21 mmol, 1 eq.) was dissolved in $H_2O$ / methanol solution (1/1, v/v, 45 mL) and heated up to 71°C. After reaching this temperature, TEA (11.2 mL, 80 mmol, 7 eq., Sigma–Aldrich) was added and the whole mixture was stirred for another 4 h. The course of the reaction was controlled by RP–HPLC and low resolution MS. The solvent was evaporated under reduced pressure and then dried in a desiccator. The product was obtained as a white solid.

d. 2–N,2–N–dimethylguanosine 5’–monophosphate **(4)**

Compound **(3)** (650 mg, 2.09 mmol, 1 eq.) was suspended in trimethyl phosphate ($PO(OCH_3)_3$, 4.4 mL, 37.5 mmol, 18 eq., Sigma–Aldrich), then pre–distilled phosphoryl chloride ($POCl_3$, 0.58 mL, 6.26 mmol, 3 eq., Merck) was added and the flask was closed with a septa. The whole reaction was stirred for 2 h on ice and the reaction was controlled by RP–HPLC. The reaction was terminated by adding 45 mL of 0.6M TEAB solution. The compound was purified on an ion exchange column using a linear gradient of TEAB buffer (0 – 0.7 M). After evaporation of the buffer, the precipitate was dried in a desiccator. Finally, the white solid was obtained.

e. 2–N,2–N–dimethylguanosine 5’–monophosphate P–imidazolide **(5)**

Compound **(4)** (0.9 g, 2.31 mmol, 1 eq.), imidazole (1.6 g, 23.1 mmol, 10 eq., Merck) and DTDP (1.6 g, 6.93 mmol, 3 eq., Sigma–Aldrich) were suspended in anhydrous DMF (7 mL). TEA (6.93 mmol, 0.97 mL, 3 eq.)

and $PPh_3$ (1.82 g, 6.93 mmol, 3 eq.), both from Sigma–Aldrich, were then added and stirred for 3 h at room temperature. After this time, the reaction was terminated by adding a solution of $LiClO_4$ (0.74 g, 6.93 mmol, 3 eq.) in cold ACN (70 mL). After cooling the mixture for 1 h in a refrigerator, the precipitate was centrifuged, washing each time with ACN. The final product was dried in a desiccator over $P_4O_{10}$ to give a light grey precipitate.

f. 2–N,2–N–dimethylguanosine 5'–diphosphate **(6)**

Compound **(5)** (954 mg, 2.16 mmol, 1 eq.) was suspended in DMF (25 mL) followed by the addition of orthophosphoric acid (V) TEA salt (3.7 g, 9.24 mmol, 4 eq.) and stirred until dissolution. $ZnCl_2$ (5.04 mg, 36.96 mmol, 16 eq., Sigma–Aldrich) was then added and stirred vigorously at room temperature for 3 h. The progress of the reaction was controlled by RP–HPLC and low resolution MS. After 3 h, the reaction was terminated by adding an aqueous solution (250 mL) of disodium EDTA (13.76 g, 36.96 mmol, 8 eq.) and $NaHCO_3$ (1/2 m EDTA) to adjust to pH 7. Compound **(6)** was purified on a DEAE Sephadex A–25 ion exchange column using a linear gradient of TEAB (0 – 0.9 M) and then evaporated to dryness and lyophilized. The product was obtained as a white powder.

g. 2–N,2–N,7–N–trimethylguanosine 5'–diphosphate **(7)**

Compound **(6)** (348.54 mg, 0.74 mmol) was dissolved in water (7.22 mL). Using glacial $CH_3COOH$, the solution was adjusted to pH 4 and the mixture was stirred for 1 h at room temperature. The progress of the reaction was controlled by RP–HPLC. Then, $(CH_3O)_2SO_2$ (0.72 mL, 7.6 µmol) was added and stirred vigorously. The solution of 1M NaOH was used to maintain pH 4. The reaction was terminated after 2.5 h and then extracted 3 times with DCM. The aqueous layer was collected and concentrated on a rotary evaporator. Compound **(7)** was purified on a DEAE Sephadex A–25 ion exchange column using a linear gradient of TEAB (0 – 0.9 M) and then evaporated to dryness and lyophilized. The product was obtained as a white powder.

h. Solid–phase synthesis of pApG **(8)**

Automatic solid phase syntheses of dinucleotide **8** was carried out using the phosphoramidite chemistry. The previously reported procedure (Mlynarska–Cieslak et al.) was applied, increasing the reaction scale to 50 µM. Modifications were introduced at the stage of cleavage of the dinucleotide from the support, increasing the time of incubation of the sample with AMA (ammonium hydroxide/methyl amine, 1/1, v/v) (1.5 h instead of 1 h) and increasing the incubation temperature (55°C instead of 37 °C). This maximised the final efficiency of this step. The further treatments were unchanged. Compound **8** was obtained as a pale grey solids with 82% of yield.

HRMS ESI (–) calcd. m/z $[M–H]^-$ $C_{20}H_{25}N_{10}O_{14}P_2^-$: 691.10324, found: 691.10381.

i. Activation of pApG (Im–pApG) **(9)**

Compound **(8)** as a triethylammonium salt (29.95 mg, 0.038 mmol, 1 eq.), imidazole (41.13 mg, 0.6 mmol, 16 eq., Merck) and DTDP (49.90 mg, 0.23 mmol, 6 eq., Sigma–Aldrich) were suspended in anhydrous DMSO (1.15 mL). TEA (31.82 µL, 0.23 mmol, 6 eq., Sigma–Aldrich) and $PPh_3$ (59.42 mg, 0.23 mmol, 6 eq., Sigma–Aldrich) were then added and stirred for 2 h at room temperature. After this time, the reaction was terminated by adding a solution of $NaClO_4$ (46.31 mg, 0.38 mmol, 10 eq.) in cold ACN (11.50 mL). After cooling the mixture for 1 h in a refrigerator, the precipitate was centrifuged, washing each time with cold ACN. The final product was dried overnight in a desiccator over $P_4O_{10}$ to give a light yellow precipitate in the yield of 91%.

j. $m_3{}^{2,2,7}$GpppApG **(10)**

Compound **(9)** as a sodium salt (141.03 mg, 0.20 mmol, 1 eq.) was dissolved in DMSO (11 mL) and compound **(7)** as triethyl ammonium salt (148.62 mg, 0.31 mmol, 1.5 eq.) was added. Dry $MgCl_2$ (222.22 mg, 1.63 mmol, 8 eq., Sigma–Aldrich) was then added and the mixture was vigorously stirred at room temperature. The reaction was controlled by performing analyses on RP–HPLC every half hour. After 5 h, the reaction was terminated by adding an aqueous solution (110 mL) of disodium EDTA (606.51 mg, 1.63 mmol, 8 eq.) and $NaHCO_3$ (1/2 m EDTA) to adjust to pH 7. The compound **(10)** was purified using Flash system followed by RP HPLC. The final synthesis product was lyophilized 3 times and its structure was confirmed by high resolution MS.

HRMS ESI (–) calcd. m/z $[M–H]^-$ $C_{33}H_{44}N_{15}O_{24}P_4^-$: 1158.16396, found: 1158.16535.

$m_3{}^{2,2,7}$GpppApG (TMG)

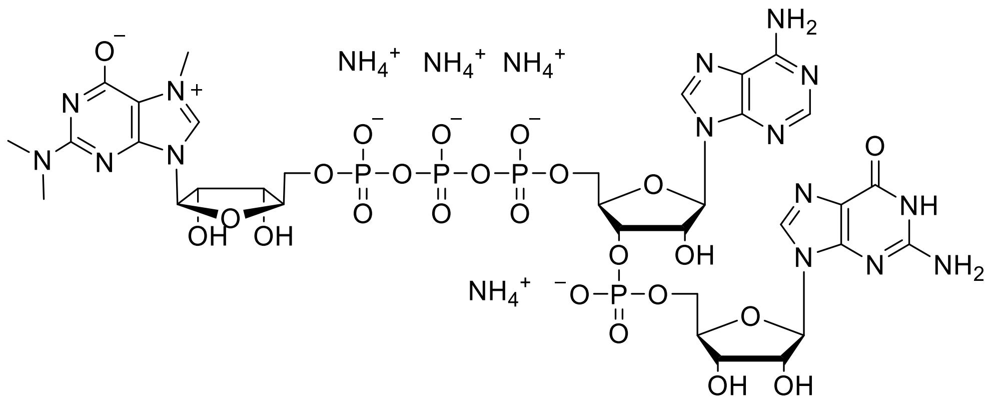


RP HPLC

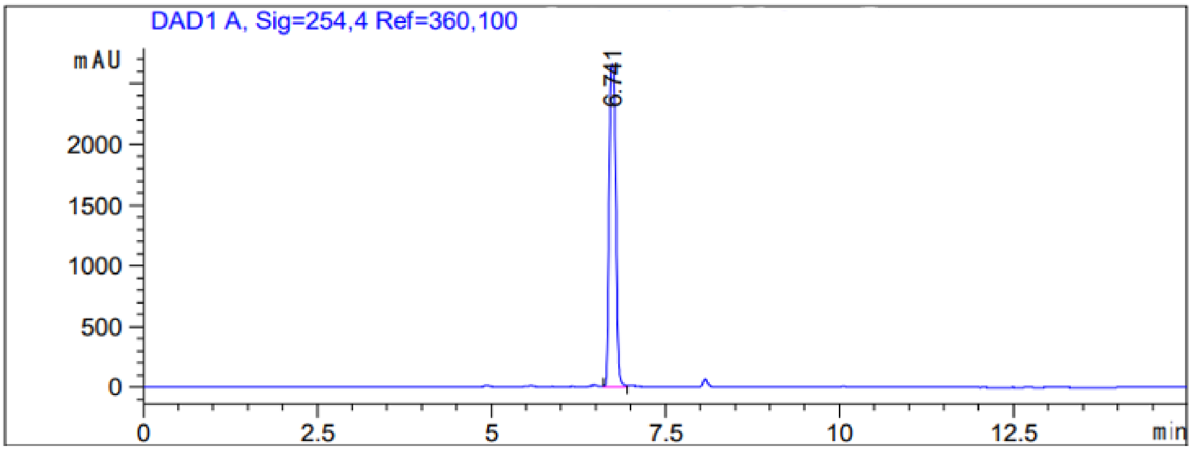

HRMS (-) ESI

(Calc. [M-H]$^-$ $C_{33}H_{44}N_{15}O_{24}P_4^-$ 1158.16396)

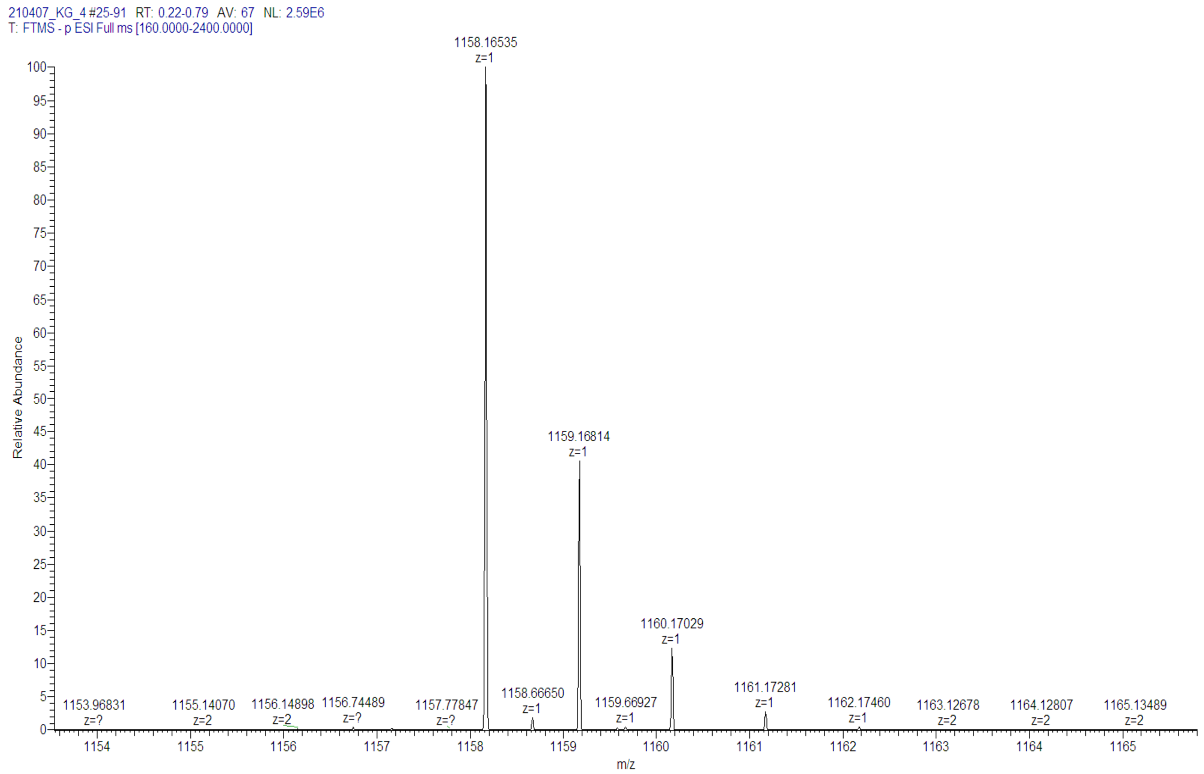


[1] Murray, M. F., Nghiem, M., and Srinivasan, A. (1995) HIV infection decreases intracellular nicotinamide adenine dinucleotide [NAD], *Biochem Biophys Res Commun 212*, 126-131.

[2] Murray, M. F., and Srinivasan, A. (1995) Nicotinamide inhibits HIV-1 in both acute and chronic in vitro infection, *Biochem Biophys Res Commun 210*, 954-959.

[3] Frindert, J., Zhang, Y., Nübel, G., Kahloon, M., Kolmar, L., Hotz-Wagenblatt, A., Burhenne, J., Haefeli, W. E., and Jäschke, A. (2018) Identification, Biosynthesis, and Decapping of NAD-Capped RNAs in B. subtilis, *Cell Reports 24*, 1890-1901.e1898.

[4] Winz, M.-L., Cahová, H., Nübel, G., Frindert, J., Höfer, K., and Jäschke, A. (2016) Capture and sequencing of NAD-capped RNA sequences with NAD captureSeq, *Nature Protocols 12*, 122.

[5] Ocwieja, K. E., Sherrill-Mix, S., Mukherjee, R., Custers-Allen, R., David, P., Brown, M., Wang, S., Link, D. R., Olson, J., Travers, K., Schadt, E., and Bushman, F. D. (2012) Dynamic regulation of HIV-1 mRNA populations analyzed by single-molecule enrichment and long-read sequencing, *Nucleic Acids Research 40*, 10345-10355.

[6] Sertznig, H., Hillebrand, F., Erkelenz, S., Schaal, H., and Widera, M. (2018) Behind the scenes of HIV-1 replication: Alternative splicing as the dependency factor on the quiet, *Virology 516*, 176-188.

[7] Huang, F. (2003) Efficient incorporation of CoA, NAD and FAD into RNA by in vitro transcription, *Nucleic Acids Research 31*, e8-e8.

[8] Mlynarska-Cieslak, A., Depaix, A., Grudzien-Nogalska, E., Sikorski, P. J., Warminski, M., Kiledjian, M., Jemielity, J., and Kowalska, J. (2018) Nicotinamide-Containing Di- and Trinucleotides as Chemical Tools for Studies of NAD-Capped RNAs, *Organic Letters 20*, 7650-7655.

[9] Honcharenko, M., Zytek, M., Bestas, B., Moreno, P., Jemielity, J., Darzynkiewicz, E., Smith, C. I., and Strömberg, R. (2013) Synthesis and evaluation of stability of m3G-CAP analogues in serum-supplemented medium and cytosolic extract, *Bioorg Med Chem 21*, 7921-7928.
